# MetaTrass: High-quality metagenome assembly on the human gut microbiome by co-barcoding sequencing reads

**DOI:** 10.1101/2021.09.13.459686

**Authors:** Yanwei Qi, Shengqiang Gu, Yue Zhang, Lidong Guo, Mengyang Xu, Xiaofang Cheng, Ou Wang, Jianwei Chen, Xiaodong Fang, Xin Liu, Li Deng, Guangyi Fan

**Author notes:** These authors contributed equally to this work (Yanwei Qi, Shengqiang Gu). Corresponding authors (Guangyi Fan,; Li Deng,.).

## Abstract

With the development of sequencing technologies and computational analysis in metagenomics, the genetic diversity of non-conserved regions has been receiving intensive attention to unravel the human gut microbial community. However, it remains a challenge to obtain enough microbial draft genomes at a high resolution from a single sample. In this work, we presented MetaTrass with a strategy of binning first and assembling later to assemble high-quality draft genomes based on metagenomics co-barcoding reads and the public reference genomes. We applied the tool to the single tube long fragment reads datasets for four human faecal samples, and generated more high-quality draft genomes with longer contiguity and higher resolution than the common combination strategies of genome assembling and binning. A total of 178 high-quality genomes was successfully assembled by MetaTrass, but the maximum of 58 was generated by the optimal common combination strategy in our tests. These high-quality genomes paved the way for genetic diversity and lineage analysis among different samples. With the high capability of assembling high-quality genomes of metagenomics datasets, MetaTrass will facilitate the study of spatial characters and dynamics of complex microbial communities at high resolution. The open-source code of MetaTrass is available at https://github.com/BGI-Qingdao/MetaTrass.

## Introduction

Through sequencing and analyzing the DNA of microbial communities directly from the environment, metagenomics has showed important roles in advancing the study of uncultured microbiomes [1, 2]. Comprehensive databases of metagenome-assembled genomes, especially for the human gut microbiome, are massively expanded to completely understand the genomic taxonomic structure of different microbiome communities according to genetic similarity [3, 4]. The progresses in metagenomics have shed new light on the study of spatial distribution and dynamics of complex microbial communities from the human gut [5, 6].

Based on the function mining of high-quality strain-resolved genomes, it is realized that genotypic differences among strains are strongly correlated with their phenotypic differences [7, 8]. The importance of intra-species non-homologous genes have been intensively studied in the field of pathogenicity, and many new species with both pathogenic and commensal strains have been found [9–11]. Indeed, the percentage of conserved intra-species homologous genes shared between strains is as low as 40% [12], and the large part of non-conservation region is thought of as the genetic origin of phenotypic diversity. Thus, complete draft genomes from a microbiome sample at the species level will enable a more comprehensive study of intra-species genome diversity, but it is still a challenge to generate sufficient high-quality genomes from metagenomic datasets.

Most of current approaches to analyze the microbiome communities are based on high-throughput and low-cost next-generation sequencing (NGS) reads. Many highly modularized computational tools have been developed such as genome assemblers, genome binners, taxonomic binners and taxonomic profilers [13–15]. The combinations of assembling first and binning later have been commonly used to generate metagenome-assembled genomes. In these strategies, a mass of short reads from a microbial community are firstly assembled to generate longer sequences by metagenomics assemblers with the consideration of uneven coverage depths of different microbial species [16–18]. Then the assembled sequences are grouped into individual genomes by genome binners based on similar *K-mer* composition and read coverage [19–21]. As a result, draft genomes with non-conserved genes are retrieved from various microbial communities. However, it is impossible to solve the assembling problem of the long inter-species repeats by the short NGS reads, so the contiguity of the draft genomes assembled by NGS reads is still not enough long for studying the long structural variations in metagenomics.

Various sequencing technologies with long-range information accompanied by specialized computational tools are promised to overcome the problem of long repeats. Third-generation single-molecule real-time sequencing (TGS) technologies developed by Pacific Biosciences and Oxford Nanopore Technology (ONT) can produce contiguous reads with lengths up to hundreds of kb, and show great potential to generate complete genomes from both cultured and uncultured microbial communities [22–24]. With using the chromatin-level contact probability information generated by high-throughput chromosome conformation capture (Hi-C) technology, more high-quality genome bins with improved contiguity can be retrieved [25]. Additionally, the co-abundance of species in multiple samples with the common *K-mer* composition are also used to improve the capability to retrieve high-quality genome bins for NGS datasets [26]. However, there are limitations for these approaches. The high sequencing error rate in TGS long reads hampers the distinction between true variations and sequencing errors. An effective contact map with Hi-C library can only be established for a draft genome with preferable contiguity. Constructing co-abundance in multiple samples ignores the genome characteristics of a single sample and increase the sequencing cost.

The co-barcoding sequencing library [27–31], an improved short-read sequencing with long-range genomic information, can provide an alternative way to improve metagenomics analyzing. In a co-barcoding library construction, long fragments sheared from DNA samples are firstly distributed into different isolated partitions, and then short-read fragments from the long fragment in the same partition are labeled with a unique barcode sequence, finally the co-barcoded fragments are sequenced by standard short-read sequencing platforms. For different co-barcoding libraries such as BGI’s single tube long fragment reads (stLFR) library [30], 10X Genomics’ linked-reads library [32] and Illumina’s contiguity preserving transposase sequencing library [28], different technical metrics in the total barcode number and the short-read coverage of the long fragment have a great impact on their powers in the downstream analysis [33–36]. The co-barcoding correlation on the draft sequences or the assembled graph have been successfully applied to improve the contiguity of assembled genomes for both large eukaryotic genomes [37–39] and metagenomes [29, 40, 41]. All these methods are still the common combination strategy in principle, leaving the inherent problem of long repeats among species with uneven abundance unsolved in efficiently constructing high-quality draft genomes for complex microbial communities.

In this work, we introduced a pipeline of **Meta**genomics **T**axonomic **R**ead **A**ssembly of **S**ingle **S**pecies (MetaTrass) based on co-barcoding sequencing data and references. Different from the common strategies, MetaTrass was a strategy of binning first and assembling later. The co-barcoding information was used not only to improve the assemblies by implementing co-barcoding assemblies, but also to simplify the dataset before assembling using microbial references with the help of taxonomic binning. We apply MetaTrass to stLFR datasets of a mock metagenome community and four real gut microbiome communities to evaluate its capability of producing high-quality draft genomes with high contiguity and high taxonomic resolution. The results were benchmarked by comparing to the common combinations of several mainstream tools. Meanwhile, the microbiome composition and genetic diversity in the four human gut samples were quantitatively analyzed with using the high-quality draft genomes assembled by MetaTrass. We expected that the high-quality draft genomes with taxonomic information at the species level assembled by our tools would be convenient to make more extensively use to investigate various microbial communities.

## Materials and methods

### Datasets

A mock microbial and four gut microbial communities were analyzed to evaluate the efficiency of MetaTrass. The mock microbial community (ZymoBIOMICS™ Microbial Community DNA Standard) consists of 8 isolated bacteria with the abundance of about 12% and 2 fungi with the abundance of about 2%. The four gut microbial DNA samples include faeces of three healthy volunteers and one patient with inflammatory bowel disease. The stLFR libraries were constructed according to the standard protocol [30]. The DNA samples were firstly sheared into long fragments, and then the long fragments were captured into a magnetic microbead with a unique barcode sequence. Finally, each long fragment was broken and hybridized with a unique barcode by the Tn5 transposase on the surface of the microbead. The stLFR libraries of the mock and the patient sample were sequenced on BGISEQ500 platform, and those of healthy samples were sequenced on MGISEQ2000 platform. The read length in the read pair was 100 bp for all datasets. The mock and three healthy sample libraries were individually allocated to a half lane, and a total of about 50 Gb raw reads were generated. The patient library was allocated to a full lane, generating about 100 Gb raw reads. Barcode sequences were extracted from the end of read2 and then replaced by numerical symbols in the read names in the FASTQ file with an in-house script. SOAPfilter_v2.2 with parameters (*-y -F CTGTCTCTTATACACATCTTAGGAAGACAAGCACTGACGACATGA -R TCTGCTGAGTCGAGAACGTCTCTGTGAGCCAAGGAGTTGCTCTGG -p -M 2 -f -1 -Q 10*) was used to clean out low-quality raw reads with adaptors, excessive confused bases, and high duplications. Finally, 55.65 Gb clean data were retained for the mock microbiome, 34.48 Gb for the first healthy sample (H_Gut_Meta01), 35.33 Gb for the second (H_Gut_Meta02), 37.88 Gb for the third (H_Gut_Meta03), and 97.20 Gb for the patient sample (P_Gut_Meta01).

### Taxonomic binning

We adopted Kraken2 (version 2.0.9-beta) [42] to classify stLFR reads into different species. Firstly, a customized reference databases were constructed according to the microbial community. Specially, references attached to the ZYMO product were used for the mock sample. The Kraken2 database of the Unified Human Gastrointestinal Genomes (UHGG) collection [3] was used to study the gut samples, and which included 4542 representative genomes at the species level. Then, the corresponding stLFR reads were classified with default parameters.

### Co-barcoding reads refining

Since a taxonomic tree of references was established to reduce the number of multiple hits of a *K-mer* from inter-species homologous sequences in Kraken2, the reads from these regions were classified into the lowest common ancient (LCA) rank higher than its corresponding species. Several works tried to reallocate these reads to species by statistical inferences using the coverage depth or co-barcoding information of intra-species homologous region of a species [43, 44]. In MetaTrass pipeline, the co-barcoding correlation between reads classified into a species and those classified into high LCA ranks was used to reduce the false negative of reads classified into high LCA rank. Reads classified into a species level is defined as the taxonomic reads of the species. In this step, we collected and refined reads for each barcode according to the number of reads in the taxonomic reads (*Num_T*) and the ratio of these reads to the total reads (*Ratio_T*). Barcodes appearing in the taxonomic reads were firstly extracted as candidates. Then, we ranked candidates first in order of *Num_T* from large to small, and then *Ratio_T* for those with the same *Num_T*. Finally, reads with a barcode of *Ratio_T* larger than a threshold were chose based on the barcode rank. Since sufficient read coverage is required for assembling a complete genome, only the read sets of one species with abundance higher than 10× were refined by co-barcoding information. The abundance of each species was roughly calculated according to the coverage depth of the taxonomic reads on the reference. Meanwhile, we set a data size threshold of the refined reads to reduce the computational consumption for species with extremely high abundance (e.g., 300×). Paired-end reads were extracted by Seqtk (version 1.3-r114-dirty) according to the barcode-related read names from the FASTQ file of clean reads. Note that there were still some false positive reads, although *Ratio_T* was set to reduce them caused by the collision of long fragments from different species in the same microbead. Sequences assembled by these reads would be further filtered as following description in the section of sequences purifying.

### Co-barcoding reads assembling

Reads of a single species with abundance higher than 10× were assembled by Supernova (version 2.1.1), which is a co-barcoding *de novo* assembler for single large eukaryotic genomes with high performances. Supernova was designed for linked-reads of 10X Genomics, which have different barcode sequences and formats from stLFR reads. Thus, we converted the stLFR reads into linked-reads FASTQ files with an in-house script. Additionally, the parameter *--accept-extreme-coverage* was set to *yes* to adapt to large coverage depth differences.

### Sequences purifying

The similarity between whole genomes based on the alignment fraction (AF) and average nucleotide identity (ANI) have been commonly adopted to circumscribe species [3, 4]. MetaTrass also used the parameters of AF and ANI between assembled contigs and the reference to purify the sequences assembled by the refined co-barcoding reads. ANI was calculated independently for each alignment. AF was defined as the ratio of total alignment length to the total contig length. The alignments with ANI larger than a threshold were counted. In our practice, we set ANI threshold to 90%, and AF threshold to 50%. The alignments between contigs and references were generated by QUAST (version 5.0.2) [45] with default parameters, except that the identity threshold to obtain valid alignment was set to 90%.

### Combinations of assembling first and binning later

In a standard analysis of NGS metagenomics dataset, the combination of *de novo* genome assembling first and binning later was commonly adopted. We compared different combinations to MetaTrass by analyzing the mock and four gut samples. In our tests, the stLFR co-barcoding reads were assembled by NGS assemblers including IDBA-UD (version 1.1.3), MEGAHIT (version 1.1.3), and MetaSPAdes (version 3.10.1) or co-barcoding assemblers including Supernova [37], Athena (version 1.3.0) [29], and CloudSPAdes (version 3.13.1) [40]. Then, all these draft assemblies were binned by two genome binners, MetaBAT2 (version 2.12.1)[21] and Maxbin2.0 (version 2.2.5) [20]. Since CloudSPAdes and Athena were not designed for stLFR reads, we made an appropriate format conversion with an in-house script where LongRanger (version 2.2.2) [46] was used. In genome assembling, Supernova was run with the same parameters as those have been adopted in MetaTrass. IDBA-UD, MEGAHIT, MetaSPAdes, Athena, and CloudSPAdes were run with default parameters. All the assembling results were deposited into CNGB Sequence Archive (CNSA) [47] (https://db.cngb.org/cnsa/) of China National GeneBank DataBase (CNGBdb) [48] with accession number CNP0002163. In genome binning, MetaBAT2 and Maxbin2.0 were run with default parameters.

### Evaluations

Both reference-based and reference-free assessments were used to evaluate the quality of assemblies obtained using different strategies. For the mock microbial community with definite references, the reference-based tool QUAST was used to evaluate contiguity and accuracy of metagenomics assemblies. Minimap2 was used to map assemblies to references and get valid alignments with the identity threshold of 95%. Then, the statistics such as genome fraction, NG50/NGA50, and number of misassemblies were assessed from the alignments with default parameters. For the real gut microbial communities, the reference-free tool CheckM (version 1.1.2) [49] were run with default parameters to evaluate the completeness and contamination of each genome from metagenomics assemblies in addition to QUAST. Following the guidance proposed in CheckM, we defined a high-quality assembly if it has >90% completeness and <5% contamination and a medium-quality assembly if it has >50% completeness and <10% contamination and does not meet the high-quality criterion. In addition, the statistics of each genome such as N50, genome size, and taxonomic rank were also obtained by CheckM, where the taxonomic rank was used to demonstrate the resolution of a genome bin.

### Variation and phylogenetic analysis

All the high-quality genomes assembled by MetaTrass were used to call variations for the four gut samples. We aligned each genome to the corresponding reference using minimap2 (2.17-r974-dirty) with parameters (*-x asm5*) to prevent an alignment extending to regions with diversity >5%. SAMtools (version 1.9) [50] and PAFtools were used to convert the BAM file of initial unsorted alignments into a PAF file of sorted alignments. We identified variations using the “call” module in PAFtools with parameters (*-L 10000*) to filter out the alignments shorter than 10,000 bp. SNVs only referred to single nucleotide substitutions, excluded single-base insertions or deletions. Insertions or deletions with length shorter than 50 bp were defined as small indels, and the others were large indels. In determination of shared variations among species in different samples, the position and sequence information of a variation were used. When variation information is the same for species genomes in different samples, the variation was shared.

We used the “classify_wf” function of GTDB-tk (version 0.3.1) [51] to conduct taxonomic annotation of the genome bins obtained using the common strategies with default parameters. Considering the procedure of UHGG database construction [4], genome bins were assigned at the species level if the AF to the close species representative genomes was higher than 30% and ANI was higher than 95%. We used FastTree (version 2.1.10) [52] to build maximum-likelihood phylogenetic trees of the high-quality genomes assembled by MetaTrass. The input of protein sequence alignments was produced by GTDB-Tk using marker gene set of 120 bacteria and 122 archaea. Interactive Tree of Life (iTOL version 4.4.2) [53] was used to visualize and annotate trees.

## Results and discussion

### MetaTrass pipeline

In this work, we developed a metagenomics assembling pipeline named MetaTrass to combine the references and long-range co-barcoding information of stLFR library. From the flowchart (**Figure 1**a), the taxonomic binning of stLFR reads were processed before the genome assembling, and different from the previous common combination strategies of assembling first and binning later. In taxonomic binning, the metagenomics stLFR reads were classified into different taxonomic ranks by Kraken2 [42]. Since the phylogenetic relations among references were used in Kraken2, only the reads from intra-species homologous region of a sample genome can be classified into the target species, but the reads from inter-species homologous and intra-species non-homologous regions were not classified effectively (Figure 1b). The reads from inter-species homologous regions were classified into the higher ranks of the target species and those from intra-species non-homologous region were unclassified or classified into irrelevant ranks. Totally, about 10% of the reads were classified into the high ranks for the four human gut datasets and about 9% of the reads were unclassified (Table S1). In co-barcoding refining, the co-barcoding correlation between the reads from intra-species homologous region and those from intra-species non-homologous and inter-species homologous region was used to refine the final reads set for a target species (Figure 1b). The barcodes of the intra-species homologous reads were firstly extracted as the candidate barcodes. Then, the final barcodes were collected by a constraint of data size and the quality of co-barcoding information. Finally, the reads with a barcode belong to the final barcodes were gathered to form the refined reads set for the target species. The constraint of data size was set to reduce computational consumption for the species with extremely high abundance. Since the barcodes with more reads classified into the target species are more possible to retain the long-range genomic information, the quality of co-barcoding information of a barcode was quantified by the number of reads classified into the species and the number ratio of these reads to total reads. In co-barcoding assembling, the refined reads of each species were independently assembled by Supernova. In practice, multiple long fragments from different species would share the same barcode in real stLFR libraries (Figure S1). Thus, the impure sequences assembled by the false positive reads from non-target species should be removed finally according to AF and ANI values of alignments between the assembly and references. Overall, the comprehensive use of co-barcoding information and references in our approach could reduce the false negative effects of taxonomic binning and the false positive effects of co-barcoding read refining.

**Figure 1.**
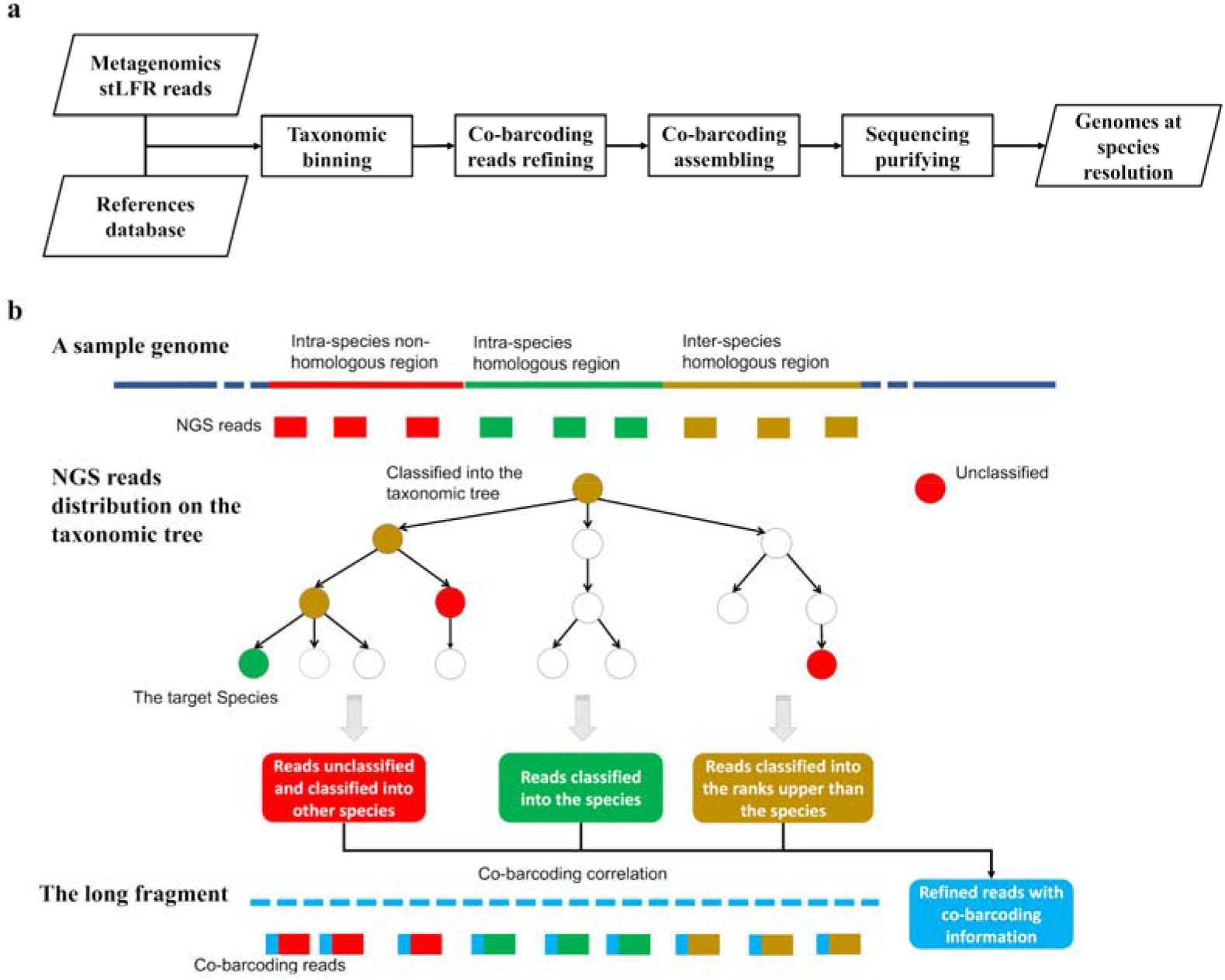
Flowchart and scheme of MetaTrass. a) Flowchart of MetaTrass assembling pipeline. b) Scheme of the homologous relation and co-barcoding correlation of difference reads sets classified by taxonomic binning.

### Assembly of the mock microbiome

The strategy of binning first and assembling later have been widely adopted to assemble haplotype genomes for eukaryotes with large sizes [54, 34]. But it has been rarely used to assemble metagenomes. We firstly applied MetaTrass to assemble stLFR read sets of the mock microbial community. Totally, up to 99.4% of reads were assigned to different datasets of species due to the simplicity of the microbial community with low inter-species homology and intra-species non-homology (Table S2). To investigate the efficiency of our strategy, we compared it with the mainstream mixed assembling strategies (Figure 2a). Besides the MetaTrass, the stLFR reads were also directly assembled by IDBA-UD, MEGAHIT, Supernova, CloudSPAdes, and Athena in the mixed assembling. Additionally, the optimal mixed assemblies of ONT reads and Illumina NGS reads in Nicholls’s work [55] were also used to make a comparison, where the ONT result was assembled by WTDBG and the NGS result was by SPAdes. The draft genome of each species in a mixed assembly was extracted by our sequence purifying module.

**Figure 2.**
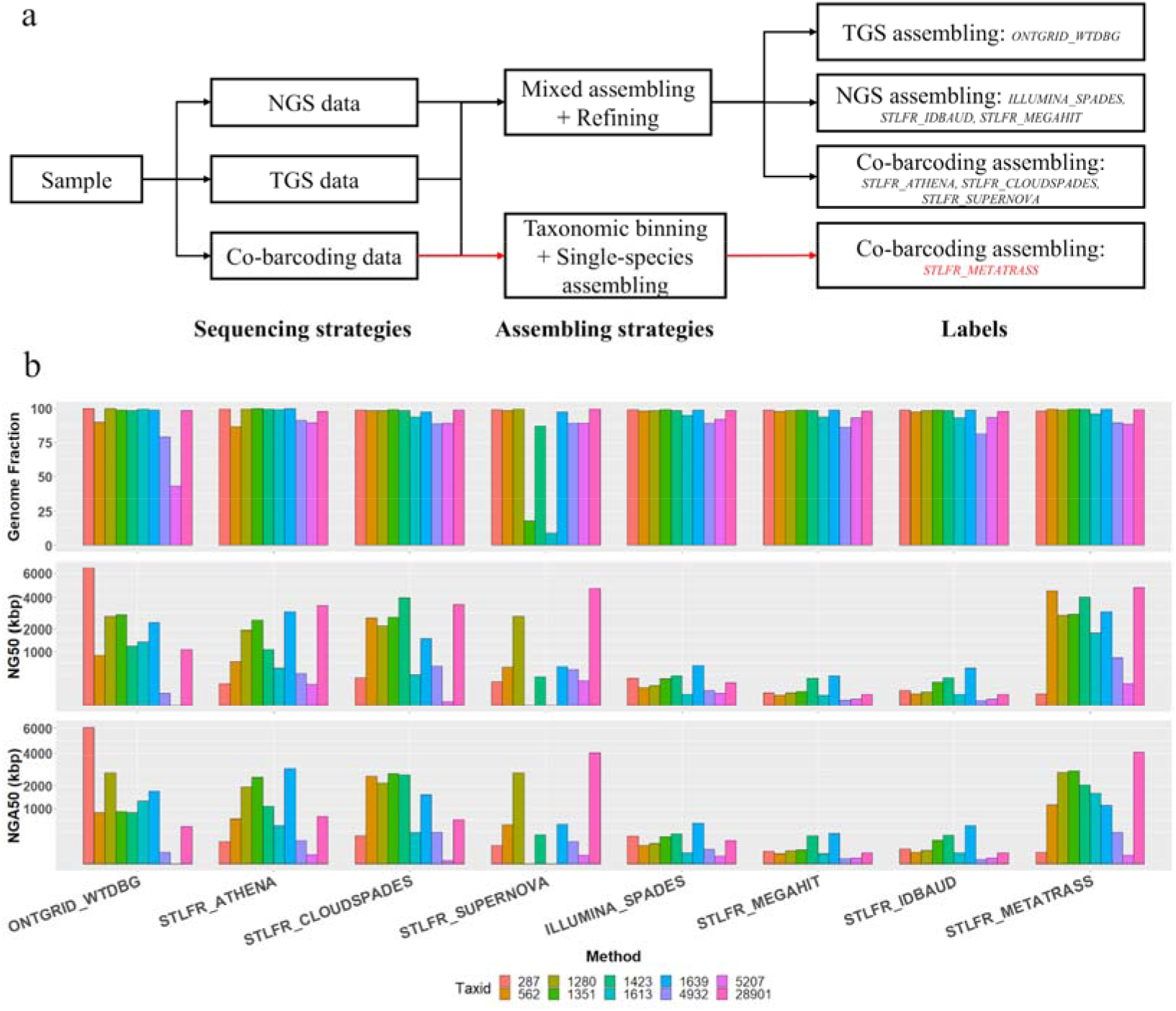
Scheme and evaluations for different strategies. a) Difference labels of the assemblies based on different sequencing and assembling strategies. b) Genome fraction, NG50 and NGA50 evaluated by QUAST for the assemblies.

Overall, our pipeline was superior in the production of draft genomes with high genome fractions and long contiguity (**Figure 2**). Two species *Enterococcus faecalis* and *Lactobacillus fermentum* were incompletely assembled by Supernova, and their genome fractions were only 17.7% and 8.9%. However, both species were properly recovered in MetaTrass, indicating that the assembling complexity caused by uneven abundances was reduced by taxonomic binning. All the assemblies by MetaTrass showed high genome fraction as those by NGS and co-barcoding assemblers designed for metagenome, which were higher than those of ONT assemblies. Compared to NGS assemblers, the co-barcoding and TGS assembler generated draft assemblies with significantly better contiguity, where Metatrass generated the best performance. MetaTrass produced seven draft genomes with NG50 around 2 Mb. Furthermore, the accuracy was guaranteed by MetaTrass, which obtained the most assemblies with NGA50 around 2 Mb. Meanwhile, assemblies by MetaTrass had less assembly errors compared to ONT assemblies (Figure S2). The average mismatch and indel numbers per 100 kb in assemblies with stLFR reads were 60 and 10, which were obviously smaller than that of the ONT assemblies.

### Assembly of four human gut microbiomes

To evaluate the robustness of our approach, we applied MetaTrass to four human faecal samples. The comprehensive genome references of UHGG were used to classify NGS reads by Kraken2, and the community compositions were estimated by the classified reads at different taxonomic ranks (Figure S3-S6). The three healthy samples had a similar microbial community, where the major microbiomes were from *Firmicutes A* phylum. This microbial community was different from the patient microbial community dominated by *Proteobacteria* which is strongly correlated with the enteric diseases caused by dysbiosis in gut microbiota [56]. The total numbers of species with higher than 10× abundance were 113, 108, 93, and 158 in H_Gut_Meta01, H_Gut_Meta02, H_Gut_Meta03, and P_Gut_Meta01 samples, respectively. The relations between these abbreviated notations and detailed sample information were descripted in the section of Materials and methods.

The genome fraction of an assembly to the reference is used to evaluate the completeness in single genome assembling. The genome fraction for all samples widely ranges from 0% to 90%, and the distributions of H_Gut_Meta01 and H_Gut_Meta02 were more concentrated than those of H_Gut_Meta03 and P_Gut_Meta01 (**Figure 3**a). However, more than half of the assembled genomes were with a genome fraction of at least 50%. Considering the large genetic diversity between sample genomes and the references [7], these results indicated that our pipeline could assemble complete genomes for species abundance of higher than 10×. The genetic diversity was also proved by the significant differences in genome fraction and the ratio of assembled length to the reference length among the four samples (Figure S7). The distributions of genomes N50 were generally dispersed, and the medians of H_Gut_Meta02 and H_Gut_Meta03 were obviously higher than those of H_Gut_Meta01 and P_Gut_Meta01 (Figure 3b). Nevertheless, the third quartiles in the box plots for the samples were larger than 100 kb, demonstrating that our pipeline had a strong capability to generate draft genomes with high contiguity. Note that for these three healthy samples plenty of ultra-long draft genomes (N50>1 Mb) was obtained, which provide possibilities to study the large genome difference in the microbiome.

**Figure 3.**
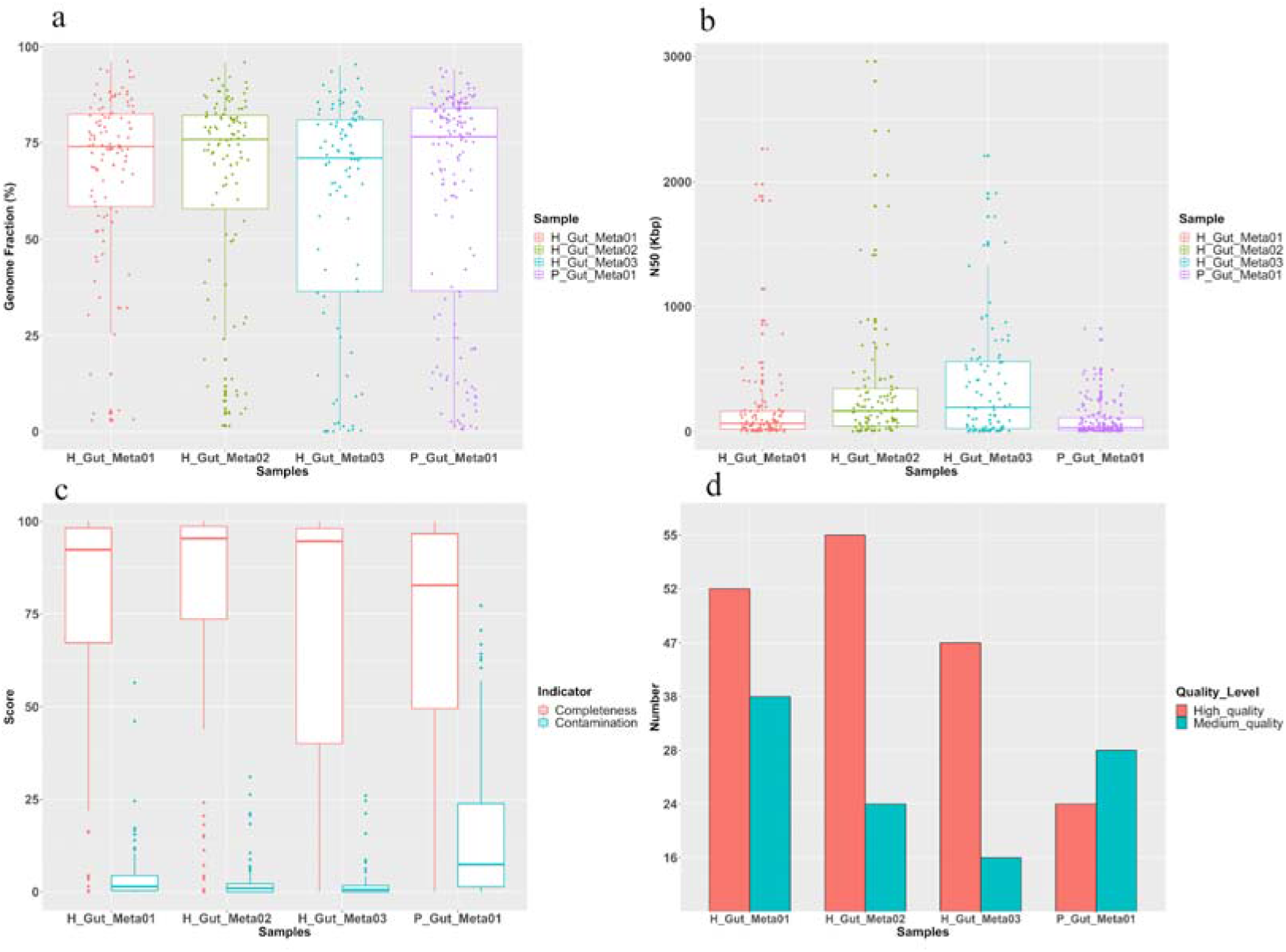
QUAST and CheckM evaluations of MetaTrass assemblies for the four human gut samples. a) Genome fraction. b) Scaffold N50. c) Box plot of completeness and contamination. d) Number of high- and medium-quality genomes.

Considering the intra-species genetic diversity, we also evaluated the quality of metagenomics assemblies based on the conserved marker genes by CheckM. The completeness medians of three healthy samples were larger than 92%, and the contamination medians were smaller than 2% (Figure 3c). The completeness of the patient sample was about 83%, and the contamination median was about 7% (Figure S8). Meanwhile, a great number of high- and medium-quality genomes were assembled by MetaTrass for the four samples (Figure 3d). 52 high-quality and 37 medium-quality genomes were produced for H_Gut_Meta01, and 55 and 24 for H_Gut_Meta02, and 47 and 16 for H_Gut_Meta03, and 24 and 28 for P_Gut_Meta01, respectively.

### Comparison to the common combination strategy

To further evaluate our approach’s efficiency, we compared it with common combinations of assembling tools and genome binning tools as listed in the section of Datasets and Methods. It should be noted that currently, there are still no genome binning tools to directly exploit the co-barcoding information. By counting the number of bins with completeness >50% and at least one conserved marker genes (Table S3), we observed that MetaTrass perform best of all these methods. Especially for P_Gut_Meta01, the optimal combination between Supernova and Maxbin2.0 obtained 66 bins with completeness higher than 50%, but it was significantly less than 117 obtained by MetaTrass.

By comprehensively analyzing the completeness, contamination and taxonomic rank of each bin, we assessed MetaTrass and common strategies in the ability to get high- and medium-quality genomes and resolution of taxonomic rank (**Figure 4**). For different samples, the best combination to produce the optimal results was different. The combinations of MetaSPAdes and Maxbin2.0, Supernova and MetaBAT2, MetaSPAdes and MetaBAT2, and Athena and MetaBAT2 is optimal for H_Gut_Meta01, H_Gut_Meta02, H_Gut_Meta03, and P_Gut_Meta01, respectively. For the four samples, the optimal results of the common strategies were still inferior to those of MetaTrass. For the example of H_Gut_meta01, the combination of MetaSPAdes and Maxbin2.0 produced 41 high- and medium-quality genomes, which was significantly less than 90 obtained by MetaTrass. There were only 3 out of totally 18 high-quality genomes with a taxonomic rank lower than the order, but 15 out of 52 for MetaTrass. Comparing the strategies only using NGS read information, the combination strategies of co-barcoding assembler and binner showed no obvious advantages in generating genomes with high quality and resolution, but MetaTrass was significantly superior to them. These results demonstrated that the usage of co-barcoding information in MetaTrass was more efficient and accurate than those in a mixed assembling.

**Figure 4.**
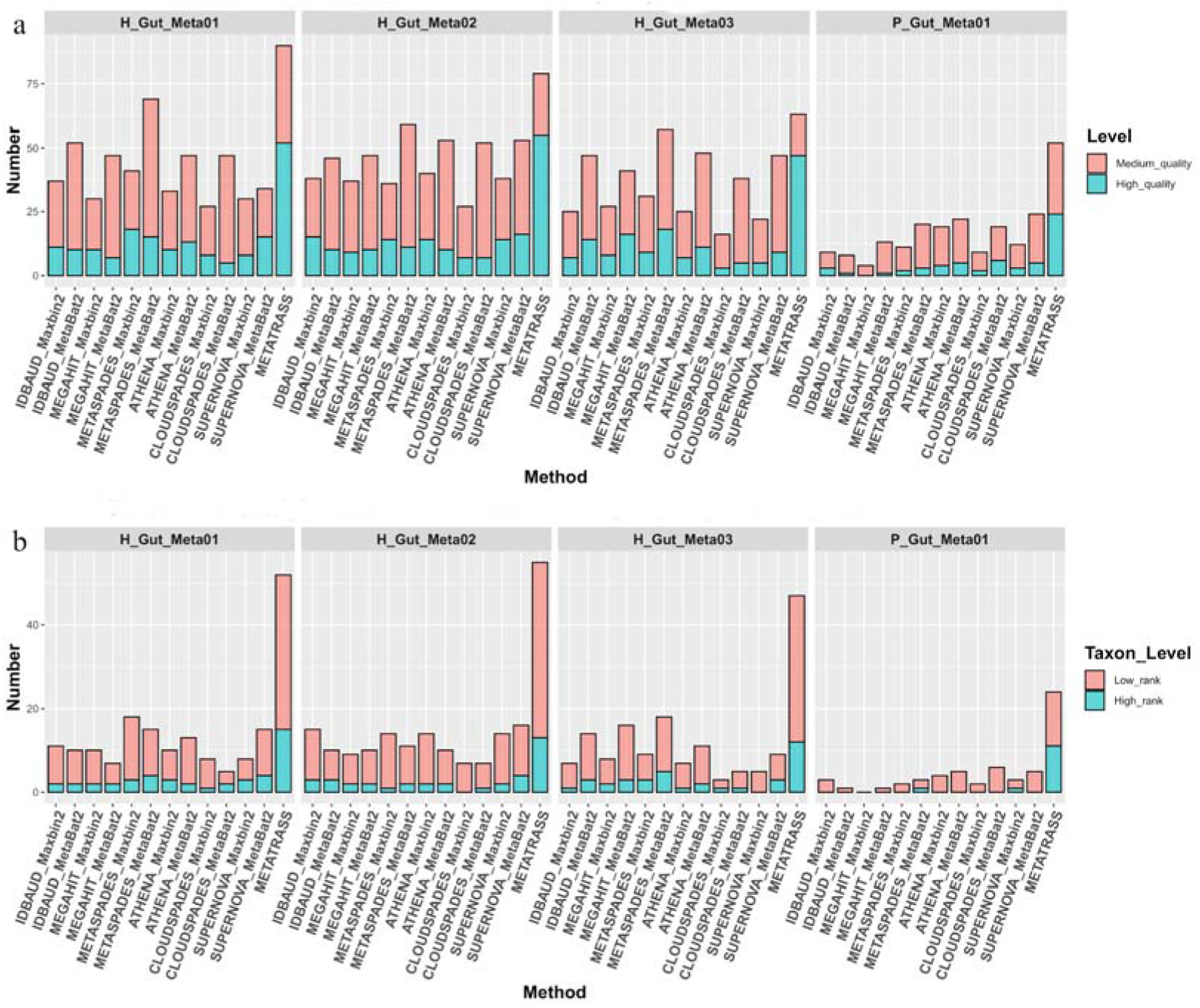
Comparison of metagenome assembling for different methods. a) Number of high- and medium-quality genomes assembled with different methods. b) Number of high-quality genomes with high- and low-rank with different methods.

The human gut microbiome composition attracts much attention due to its strong correlation with personality traits [57]. To compare the microbiome composition structures of the high-quality genomes with different methods, we uniformly classified the high-quality genome bins into species using GTDB-tk. Using the large number of high-quality genomes obtained by MetaTrass, the phylogenetic trees of these genomes were constructed and the corresponding N50 were attached in the left histogram as shown in **Figure 5**. Meanwhile, the high-quality genome bins obtained by the common strategies were marked in red in the middle heat map (Figure 5), if the genome of the same species were also assembled by MetaTrass. The topology of the phylogenetic tree of genomes assembled by MetaTrass gave comprehensive insights of the microbial composition structure. From the trees in Figure 5 and Figure S9-S11, the numbers of the order with high-quality genomes assembled by MetaTrass were 9, 11, 7, and 7 for H_Gut_Meta01, H_Gut_Meta02, H_Gut_Meta03, and P_Gut_Meta01, respectively. Notably, some orders contained more than 5 high-quality genomes, and this provide convenience to study the microbiome structure at the genome-wide scale. For the sample of H_Gut_Meta01 (Figure 5), there were 27 high-quality genomes classified into *Lachnospirales* order and 14 into *Oscillospirales*. These two were exactly the dominating orders according to the taxonomic abundance distribution. Similar results were obtained for the other two healthy samples (Figure S9 and S10), indicating that the microbiome with higher sequencing coverage could be better assembled in MetaTrass. In contrast, the orders with more than 5 high-quality genomes were *Enterobacterales* and *Actinomycetales* for P_Gut_Meta01 (Figure S11). The obvious difference between the healthy and patient samples was consistent with the microbial compositions differences observed in the taxonomic binning results. MetaTrass successfully assemble most of the high-quality genomes of all common combinations in our tests. For instance, they generated 137 genome bins, while only 25 genome bins were not assembled by MetaTrass (Figure 5). From the heat maps, most of the common strategies could assemble draft genomes for each order, but the total numbers in each order were relatively small. The maximal number of genomes in one order was 6 and obtained by the combination of Supernova and MetaBAT2 for *Lachnospirales*. Moreover, 146 of 179 high-quality genomes were with N50 values larger than 100 kb, demonstrating that MetaTrass had a strong ability to improve the contiguity of assemblies.

**Figure 5.**
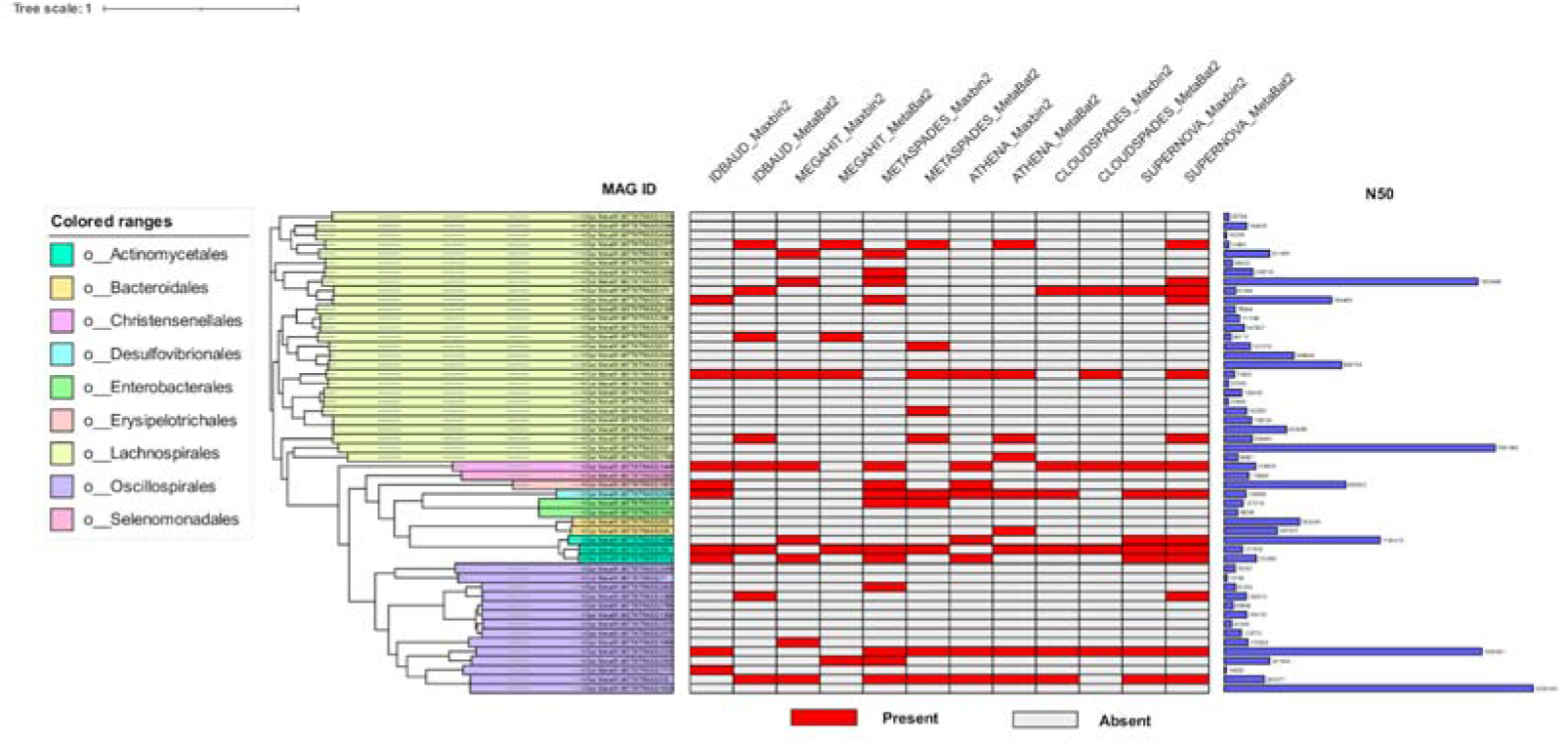
Phylogenetic tree of the high-quality genomes assembled by MetaTrass for H_Gut_Meta01. The phylogenetic tree is on the left. Distribution of the high-quality genomes assembled by other methods are colored as red in the middle heat map. N50 of each high-quality genome is shown in the right histogram.

### Genetic diversity in different samples

Different types of variations in gut microbiomes are strongly associated with host health, and the genetic diversity among different microbiomes has been intensively studied to unravel the genetic origin of phenotypic difference among people of different regions or health status [58, 59]. By aligning draft genomes to the references, we called variations for high-quality genomes for each species in different samples, including single nucleotide variations (SNV), small and large indels. For different variations, the numbers of SNV were significantly larger than those of the small and large indels for the four samples (Figure S12). Three healthy samples showed close variation numbers, which were obviously larger than those of the patient. It come from fewer alignments for the patient sample according to the QUAST evaluation. However, when we removed the effect of the total aligned length by calculating the SNV density, the patient sample showed denser SNV than the healthy samples (Figure S12d). The median was about 21 for the patient sample, but about 9 for the healthy samples. This difference could be introduced by the individual’s physiological state, which was related to the diseases and also to the territory or race [4].

Based on the taxonomic information of high-quality genomes, we found 15 species shared by three samples, where 14 species appeared in the three healthy samples but only one species of *Escherichia* appeared in the patient and two healthy samples. By analyzing the SNV density and intersection of variations between different samples for each species in three healthy samples, we further investigated the genetic diversity between species from different samples. The SNV densities were different for different species even in the same sample, but similar for the same species in different sample (**Figure 6**a). From Figure 6b to 6d, the number of unique and shared variations in different types significantly fluctuated for different species, but their difference among samples showed great consistency. The total shared numbers between H_Gut_Meta01 and H_Gut_meta02 were obviously more than those between H_GutMeta03 and the other two samples for all variations. Furthermore, the ratio of large indels shared by all three samples to the total number was much smaller than those of SNVs and small indels. These results demonstrated that large variations were more specific than small variations in the huge genetic diversity between different samples, were consistent with the observation in the study of association between host health and structural variations in gut microbiome [58].

**Figure 6.**
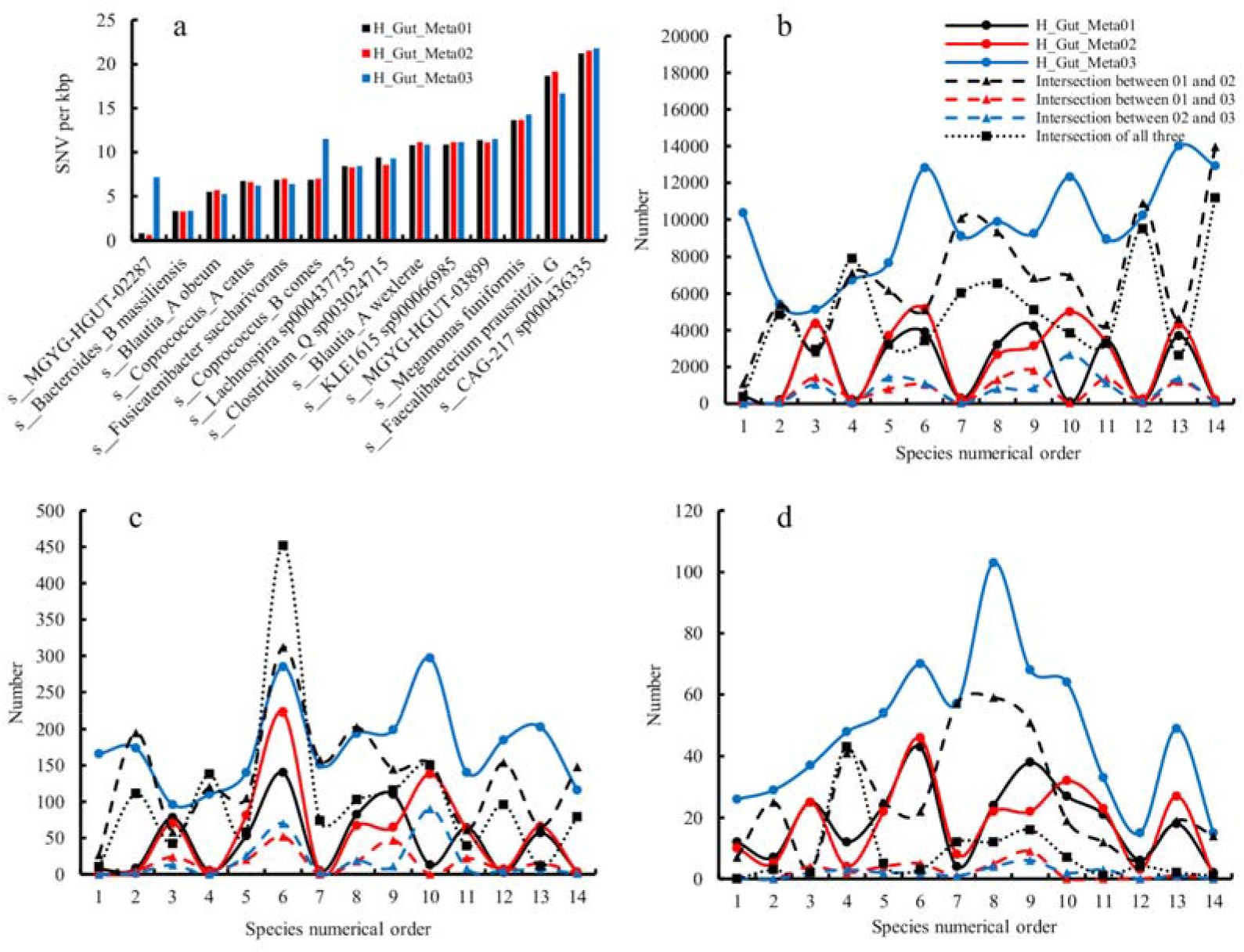
SNV density and number of unique and shared variations for each species appearing in all three healthy samples. a) is the SNV density. b), c) and d) are the number of SNVs, small and large indels, respectively. The species numerical order in b), c) and d) corresponds to the appearance order of species from left to right in a).

### Computational performance

Runtime and used thread number of each assembler were recorded for all the human gut datasets (**Table 1**). Most of the assemblers were test on 24 Intel(R) Xeon(R) Silver 4116 CPU @ 2.10GHz, except for Athena and Supernova which were test on HPC Cluster for their large memory requirements. The thread number used in each assembler was the same for different samples. The time consumption of the format conversion from stLFR reads to 10X linked-reads were not included, and was about 500 minutes for dataset with 50 Gb with one thread. We found that MetaTrass was less time consuming than Athena but more than other assemblers. This may come from that both MetaTrass and Athena contained many sub-assembling, which took most of the time among all sub-processes in MetaTrass (Table S4). Since the sub assembling was independent, it could be run in parallel to further speed up the assembling by increasing the parallel number and the parallel number was 8 in default.

**Table 1.** Runtimes and thread number of each assembler for all the human gut datasets.

| Assembler | Thread<br>number | Runtime (min) |  |  |  |
| --- | --- | --- | --- | --- | --- |
|  | All<br>samples | H_Gut_meta01 | H_Gut_Meta02 | H_Gut_Meta03 | P_Gut_Meta01 |
| IBDA-UD | 6 | 863 | 884 | 911 | 2657 |
| MEGAHIT | 16 | 179 | 161 | 163 | 611 |
| MetaSPAdes | 16 | 1478 | 1289 | 1429 | 3459 |
| CloudSPAdes | 16 | 1024 | 1163 | 1039 | 2627 |
| Supernova | 8 | 1249 | 864 | 1098 | 6776 |
| Athena | 16 | 13813 | 8689 | 6361 | -- |
| MetaTrass | 16 | 5145 | 2631 | 3147 | 8363 |

*Note:* The exact runtime of assembling P\_Gut\_Meta01 sample by Athena was not collected correctly due to several uncontrolled interrupts on HPC cluster.

## Conclusion

High-quality genomes at species level are strongly demanded to investigate the genetic origins of diseases associated with the human gut, but how to get sufficient number of them in one sample is still a challenge due to the inter-species repeats and uneven abundance in metagenomics assembling. In this work, we developed a tool to get high-quality genomes with high taxonomic resolutions by combining the co-barcoding information with public references. Compared with the common combination strategies, our pipeline generated a large number of high-quality genomes for the human microbiome co-barcoding datasets. Meanwhile, plenty of draft genomes were also assembled with NG50 values of larger than 1 Mb, some of which were even longer than the references for both mock and human gut datasets. For all the four real gut samples, 178 draft genomes with high completeness and low contamination were generated, but their genome fractions relative to the references were low. The differences between the sample genomes assembled by MetaTrass and the reference genomes demonstrated that the co-barcoding information could be used to reduce the false negative reads in taxonomic binning. These reads retrieved from inter-species homologous and intra-species non-homologous regions by co-barcoding refining could significantly improve the assembly results. For the patient sample, the number of high-quality genomes with long contiguity assembled by MetaTrass was significantly larger than that without co-barcoding refining (Figure S13).

The efficiency of our pipeline depended on the co-barcoding information quality including the read coverage and length of long fragments. By aligning reads to the species reference, we calculated the genome fraction with different read coverage depths for different read sets including the taxonomic reads, the refined reads, and all reads. According to the genome fraction with high coverage depths, we evaluate the efficiency of the co-barcoding refining. From the results of species with medium abundance in P_Gut_Meta01 (Figure S5), We observed that the fraction with high depths of the refined reads was higher than those of the taxonomic reads, but still lower than those of all aligned reads. These results indicated that there were still some false negative reads introduced by the low coverage or short length of long fragments. Thus, improvements on co-barcoding library and the co-barcoding refining would improve the performance of MetaTrass.

In summary, the application of MetaTrass in human gut samples showed great promise of generating high-quality genomes for real complex microbial community at a high resolution. With the increasing number of reference genomes from various microbial communities and the development of co-barcoding sequencing library, the combination strategy of binning first and assembling later in MetaTrass will be extended and facilitate the investigation of the association between host phenotypes and microbial genotypes for different microbial communities.

## Supporting information

Suppporting information

## Acknowledgements

This research was supported by the National Key Research and Development Program of China (2018YFD0900301-05), and Science Technology and Innovation Committee of Shenzhen Municipality of China (SGDX20190919142801722). We would thank Yufen Huang and many other BGI-Shenzhen employees for fruitful discussions in the development and performance test.

## Authors’ contributions

Li Deng, Guangyi Fan and Yanwei Qi contributed to the software design. Yanwei Qi, Shengqiang Gu, Yue Zhang and Lidong Guo contributed to the software implementation. Li Deng, Yanwei Qi, Shengqiang Gu, Mengyang Xu and Jianwei Chen contributed to data analyses. Xiaofang Chen, Ou Wang and Xiaodong Fang contribute to the data curation, collection. Guangyi Fan, Li Deng and Xin Liu contributed to the benchmarking design. All authors contributed to the manuscript writing. Li Deng and Guangyi Fan supervised the project. All authors read and approved the final manuscript.

## Data availability statement

MetaTrass is freely available at https://github.com/BGI-Qingdao/MetaTrass. The assembling results of the four human faecal samples were deposited into CNSA (https://db.cngb.org/cnsa/) of CNGBdb with accession number CNP0002163 and available from authors upon reasonable request and with permission of CNGBdb. The metagenomics stLFR datasets used in the study were available from the corresponding author on reasonable request.

## Supporting information

**Table S1** Read number on different ranks classified by Kraken2 for the four gut samples.

**Table S2** Classified read information of the mock dataset.

**Table S3** The overall view of genome bins obtained by MetaTrass and all common strategies “Comp >50%” means the completeness higher than 50%.

**Table S4** The runtime of MetaTrass step by step for all human gut datasets.

**Table S5** Genome fraction with different coverage depths for different read sets including the taxonomic read (TR), refined reads by co-barcoding (BR), and total reads (Total) for five species with medium abundances in P_Gut_Meta01.

**Figure S1** The probability of barcodes with long fragments from different species for four gut samples.

**Figure S2** Mismatches and Indels of different assemblies for the mock dataset.

**Figure S3** Distributions of classified reads at different phyla for four gut samples.

**Figure S4** Distributions of classified reads at different classes for four gut samples.

**Figure S5** Distributions of classified reads at different orders for four gut samples.

**Figure S6** Distributions of classified reads at different families for four gut samples.

**Figure S7** Genome faction and ratio of assembly length to reference length of all species assembled in MetaTrass for four gut samples, and the species are ordered by the completeness.

**Figure S8** Two-dimensional scatter plot of completeness and contamination evaluated by CheckM for four gut samples.

**Figure S9** Phylogenetic tree of the high-quality genomes assembled by MetaTrass for H_Gut_Meta02. The phylogenetic tree is on the left. Distribution of the high-quality genomes assembled by other methods are colored as red in the middle heat map. N50 of each high-quality genome is shown in the right histogram.

**Figure S10** Phylogenetic tree of the high-quality genomes assembled by MetaTrass for H_Gut_Meta03. The phylogenetic tree is on the left. Distribution of the high-quality genomes assembled by other methods are colored as red in the middle heat map. N50 of each high-quality genome is shown in the right histogram.

**Figure S11** Phylogenetic tree of the high-quality genomes assembled by MetaTrass for P_Gut_Meta01. The phylogenetic tree is on the left. N50 of each high-quality genome is shown in the right histogram. Because the genome bins obtained by the combination strategies cannot be classified into species by GTDB-tk, the heat map is not showed for this sample.

**Figure S12** Box plot of variations. Box plots of SNVs (a), small indels (b), large indels (c) and SNV density (d) called from the high-quality genomes for four gut samples.

**Figure S13** Number of genomes with different quality (a) and contiguity (b) assembled by MetaTrass and MetaTrass_TR for the patient gut sample. Since MetaTrass_TR excluded the co-barcoding refining process compared to MetaTrass, the input dataset of co-barcoding assembling in MetaTrass_TR is the taxonomic reads set. Only the high-quality genomes are considered to count the number of genomes with different contiguity.

