## Supplementary material for "MetaTrass: High-quality metagenome assembly on the human gut microbiome by co-barcoding sequencing reads": Suppporting information

**Table S1** Read number on different ranks classified by Kraken2 for the four gut samples.

|  | H_Gut_Meta01 | H_Gut_Meta02 | H_Gut_Meta03 | P_Gut_Meta01 |
| --- | --- | --- | --- | --- |
| Unclassified | 13807390 | 14713967 | 20358285 | 54591902 |
| Root | 19193 | 27560 | 12302 | 2671 |
| Domain | 3563468 | 3816148 | 3650219 | 2241393 |
| Phylum | 53639 | 4050 | 3397 | 214157 |
| Class | 2504853 | 2382908 | 2369599 | 368047 |
| Order | 1008806 | 1289630 | 1050150 | 646074 |
| Family | 5727372 | 5984160 | 5831847 | 5546802 |
| Genus | 7387356 | 6590155 | 8766559 | 53721890 |
| Species | 138332261 | 141834907 | 147366577 | 465500934 |

**Table S2** Classified read information of the mock dataset.

| Taxid | Species Name | Theoretic genomic DNA | Reference length (bp) | classified read count | Coverage depth |
| --- | --- | --- | --- | --- | --- |
| 1280 | Staphylococcus_aureus | 12 | 2730326 | 6725076 | 492.621 |
| 1351 | Enterococcus_faecalis | 12 | 2845392 | 75166780 | 5283.4 |
| 1423 | Bacillus_subtilis | 12 | 4045677 | 38992509 | 1927.61 |
| 1613 | Lactobacillus_fermentum | 12 | 1905333 | 31651503 | 3322.41 |
| 1639 | Listeria_monocytogenes | 12 | 2992342 | 22911609 | 1531.35 |
| 287 | Pseudomonas_aeruginosa | 12 | 6792330 | 4103526 | 120.828 |
| 28901 | Salmonella_enterica | 12 | 4809318 | 10133379 | 421.406 |
| 4932 | Saccharomyces_cerevisiae | 2 | 12843354 | 54904223 | 854.983 |
| 5207 | Cryptococcus_neoformans | 2 | 29176277 | 12497637 | 85.6699 |
| 562 | Escherichia_coli | 12 | 4875441 | 18123742 | 743.471 |

**Table S3** The overall view of genome bins obtained by MetaTrass and all common strategies “Comp >50%” means the completeness higher than 50%.

|  | H_Gut_meta01 | | H_Gut_Meta02 | | H_Gut_Meta03 | | P_Gut_Meta01 | |
| --- | --- | --- | --- | --- | --- | --- | --- | --- |
| Methods | Comp  >50% | Total | Comp  50% | Total | Comp  >50% | Total | Comp  >50% | Total |
| IDBA-UD+MetaBat2 | 59 | 145 | 52 | 154 | 51 | 124 | 8 | 65 |
| IDBA-UD+Maxbin2.0 | 81 | 124 | 78 | 113 | 64 | 109 | 40 | 77 |
| MEGAHIT+MetaBat2 | 50 | 134 | 52 | 132 | 43 | 115 | 14 | 81 |
| MEGAHIT+Maxbin2.0 | 81 | 117 | 76 | 111 | 59 | 94 | 37 | 67 |
| METASPADES+MetaBat2 | 76 | 169 | 65 | 157 | 59 | 135 | 26 | 113 |
| METASPADES+Maxbin2.0 | 95 | 121 | 85 | 111 | 66 | 109 | 55 | 86 |
| ATHENA+MetaBat2 | 60 | 166 | 62 | 167 | 55 | 125 | 37 | 113 |
| ATHENA+Maxbin2.0 | 77 | 103 | 65 | 91 | 49 | 74 | 62 | 82 |
| CLOUDSPADES+MetaBat2 | 53 | 188 | 60 | 156 | 42 | 166 | 24 | 116 |
| CLOUDSPADES+Maxbin2.0 | 85 | 109 | 73 | 110 | 68 | 97 | 48 | 77 |
| SUPERNOVA+MetaBat2 | 56 | 164 | 59 | 142 | 53 | 116 | 38 | 118 |
| SUPERNOVA+Maxbin2.0 | 72 | 106 | 66 | 92 | 43 | 67 | 66 | 106 |
| MetaTrass | 99 | 112 | 86 | 105 | 67 | 82 | 117 | 154 |

**Table S4** The runtime of MetaTrass step by step for all human gut datasets.

|  | H_Gut_Meta01 | H_Gut_Meta02 | H_Gut_Meta03 | P_Gut_Meta01 |
| --- | --- | --- | --- | --- |
| Taxonomic binning | 28min | 19min | 21min | 58min |
| Co-barcoding reads refining | 2h37min | 2h30min | 2h36min | 8h1min |
| Co-barcoding reads Assembling | 3d11h35min | 1d16h59min | 2d1h26min | 5d10h17min |
| Seqeuences Purifying | 3min | 3min | 4min | 7min |
| Total | 3d13h45min | 1d19h51min | 2d4h27min | 5d19h23min |

**Table S5** Genome fraction with different coverage depths for different read sets including the taxonomic read (TR), refined reads by co-barcoding (BR), and total reads (Total) for five species with medium abundances in P_Gut_Meta01.

|  | s__Veillonella parvula | | | s__Klebsiella quasivariicola | | | s__MGYG-HGUT-03797 | | | s__Streptococcus mitis_AC | | |
| --- | --- | --- | --- | --- | --- | --- | --- | --- | --- | --- | --- | --- |
| Coverage | TR | BR | Total | TR | BR | Total | TR | BR | Total | TR | BR | Total |
| >0x | 100.00 | 100.00 | 100.00 | 99.95 | 100.00 | 100.00 | 99.98 | 99.99 | 100.00 | 99.99 | 100.00 | 100.00 |
| >=4x | 96.94 | 99.39 | 99.88 | 54.64 | 98.64 | 99.51 | 95.95 | 98.70 | 99.52 | 94.54 | 97.96 | 99.24 |
| >=10x | 93.20 | 98.32 | 99.63 | 37.70 | 94.34 | 99.03 | 89.26 | 96.15 | 98.50 | 87.45 | 95.19 | 97.86 |
| >=30x | 83.68 | 94.73 | 98.52 | 27.90 | 66.67 | 98.37 | 69.49 | 87.46 | 96.04 | 70.15 | 88.60 | 95.72 |
| >=100x | 49.88 | 65.52 | 92.00 | 19.32 | 21.10 | 97.51 | 32.80 | 42.22 | 90.65 | 34.13 | 46.70 | 89.59 |

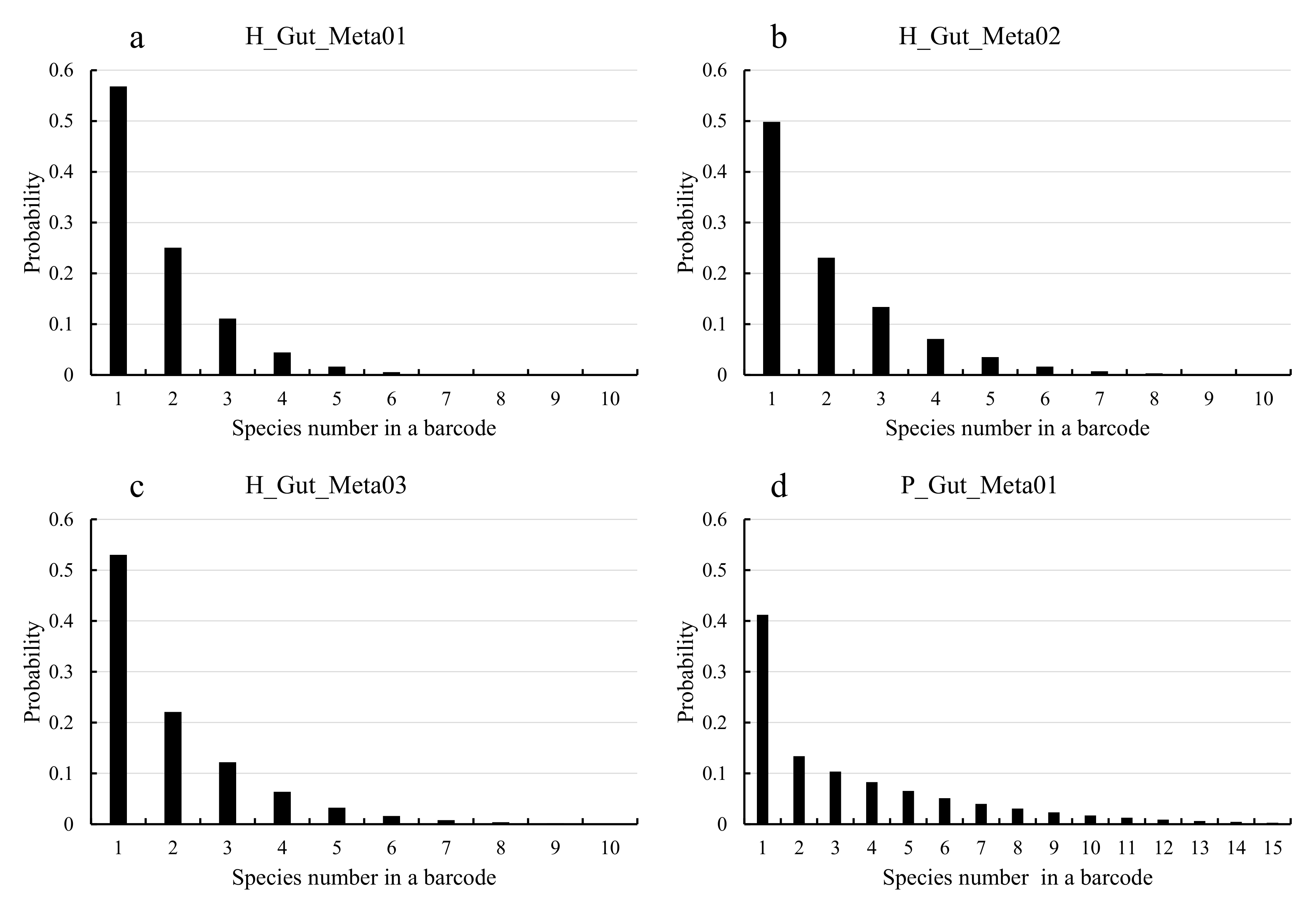

**Figure S1** The probability of barcodes with long fragments from different species for four gut samples.

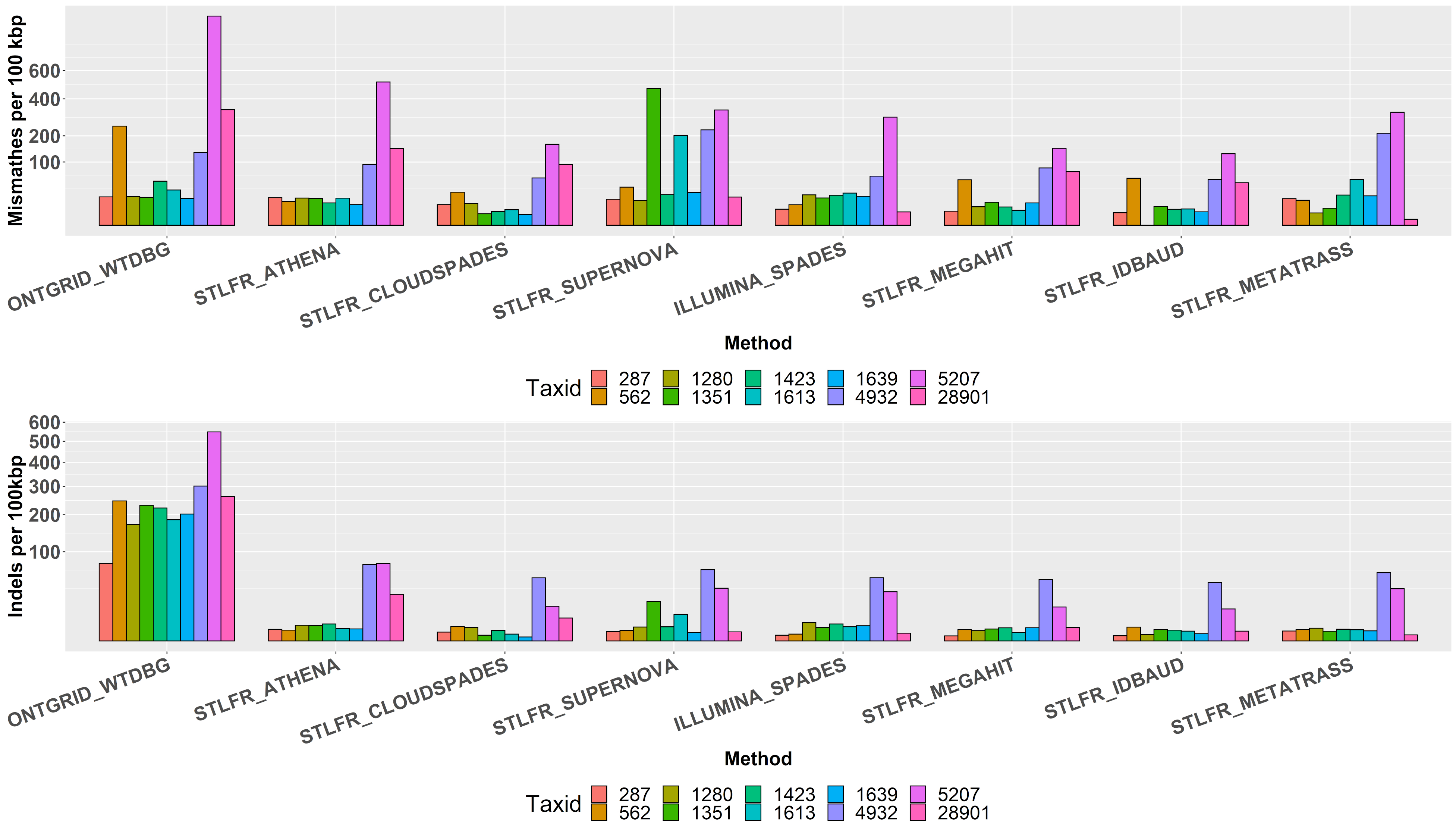

**Figure S2** Mismatches and Indels of different assemblies for the mock dataset.

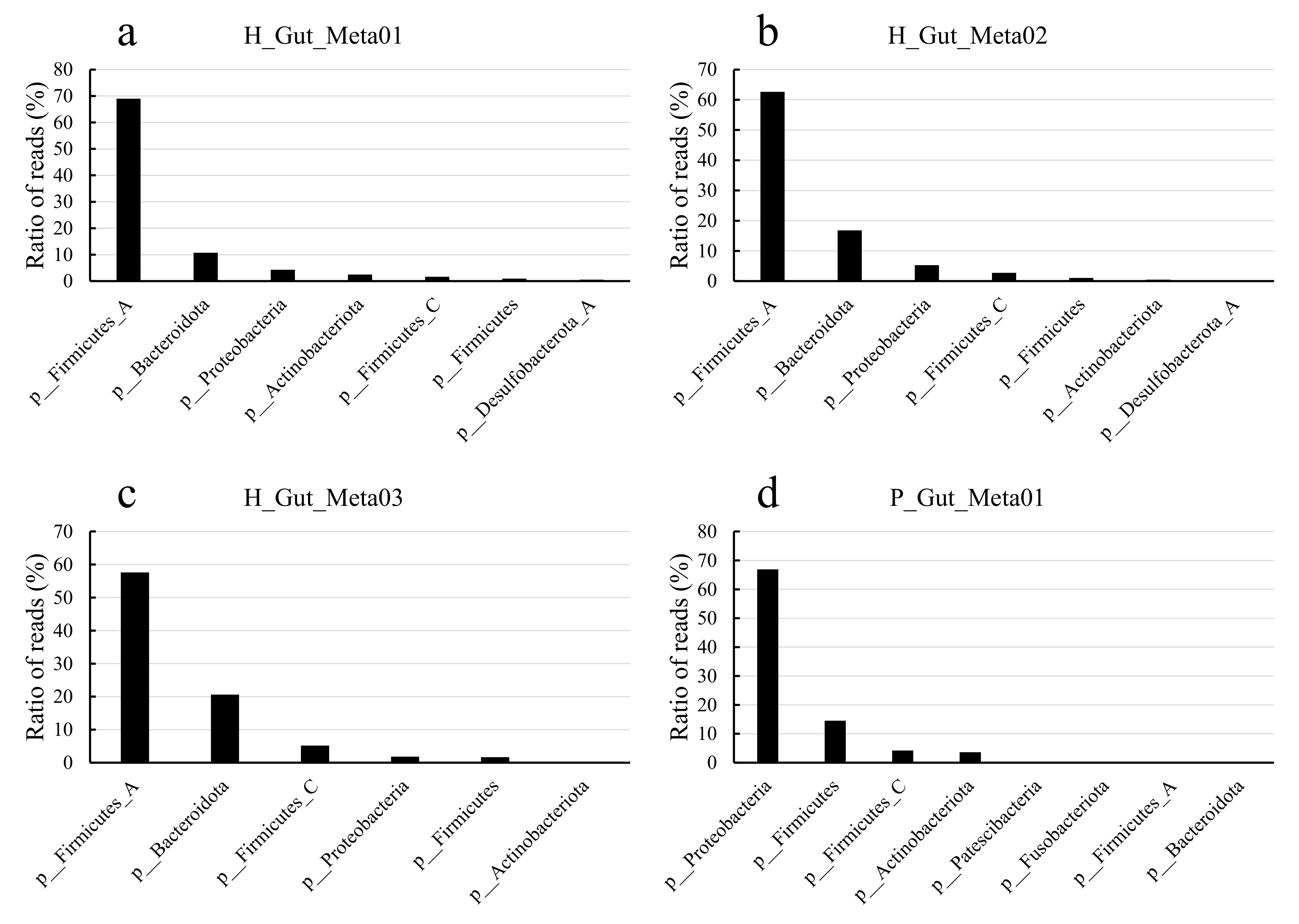

**Figure S3** Distributions of classified reads at different phyla for four gut samples.

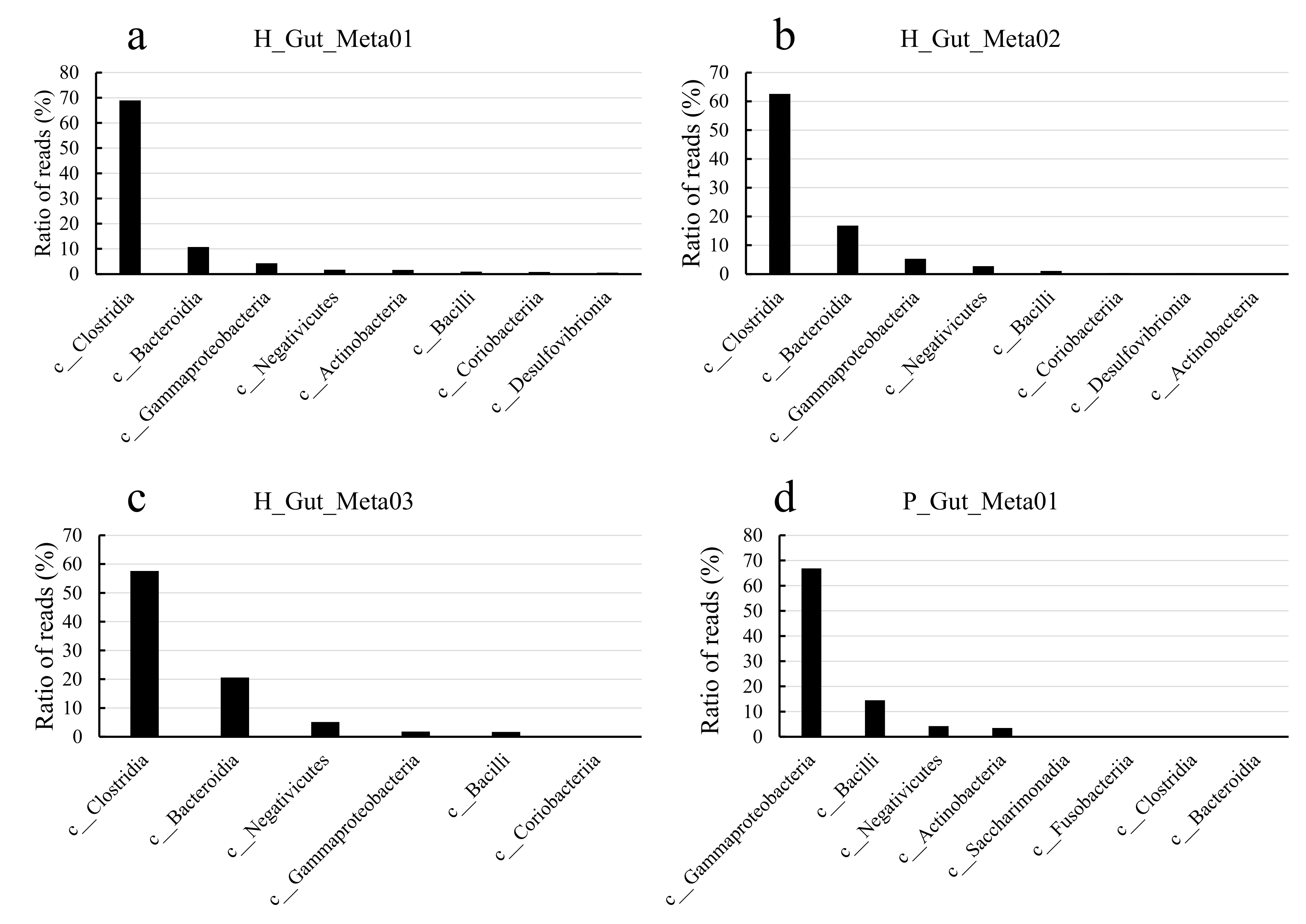

**Figure S4** Distributions of classified reads at different classes for four gut samples.

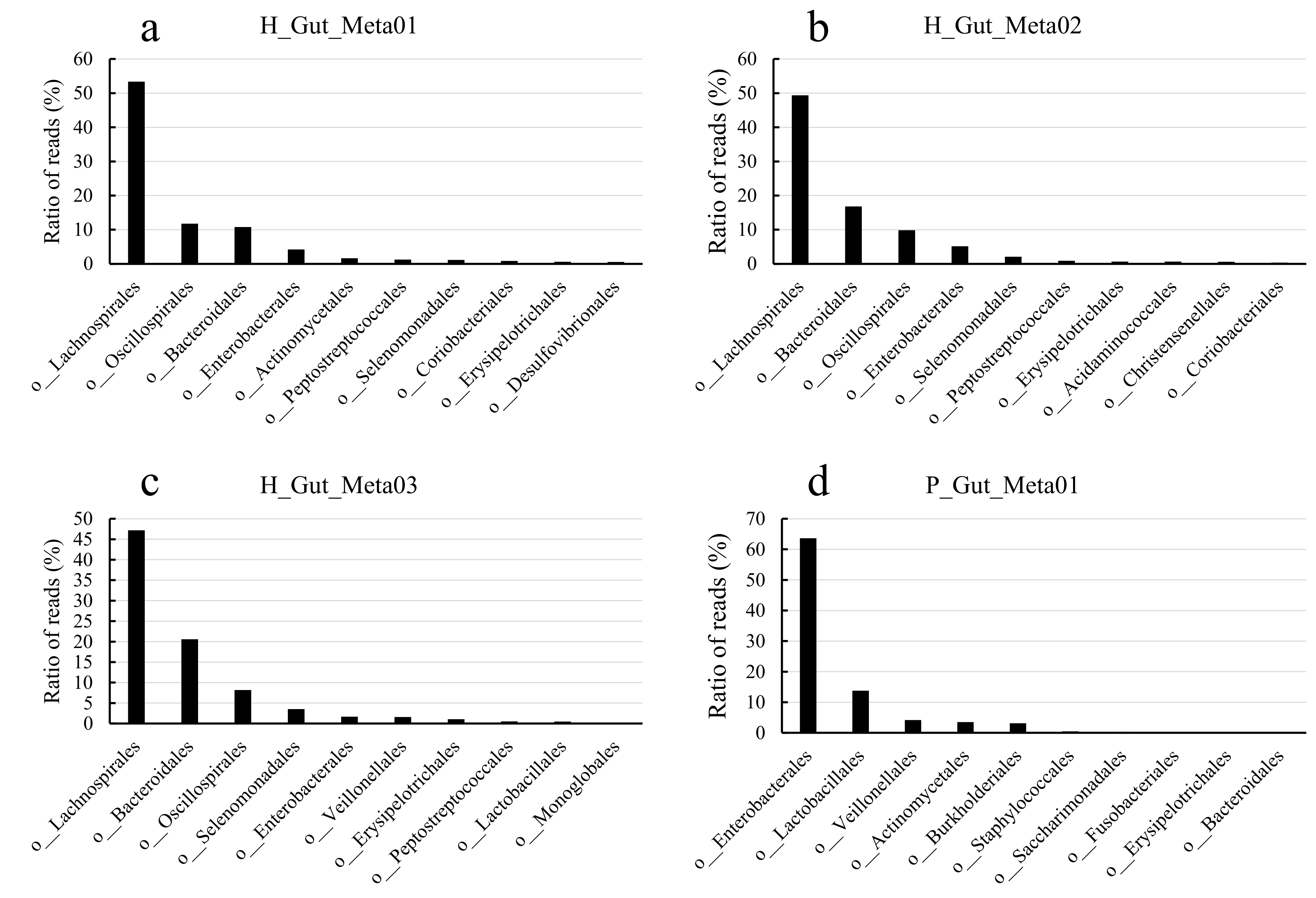

**Figure S5** Distributions of classified reads at different orders for four gut samples.

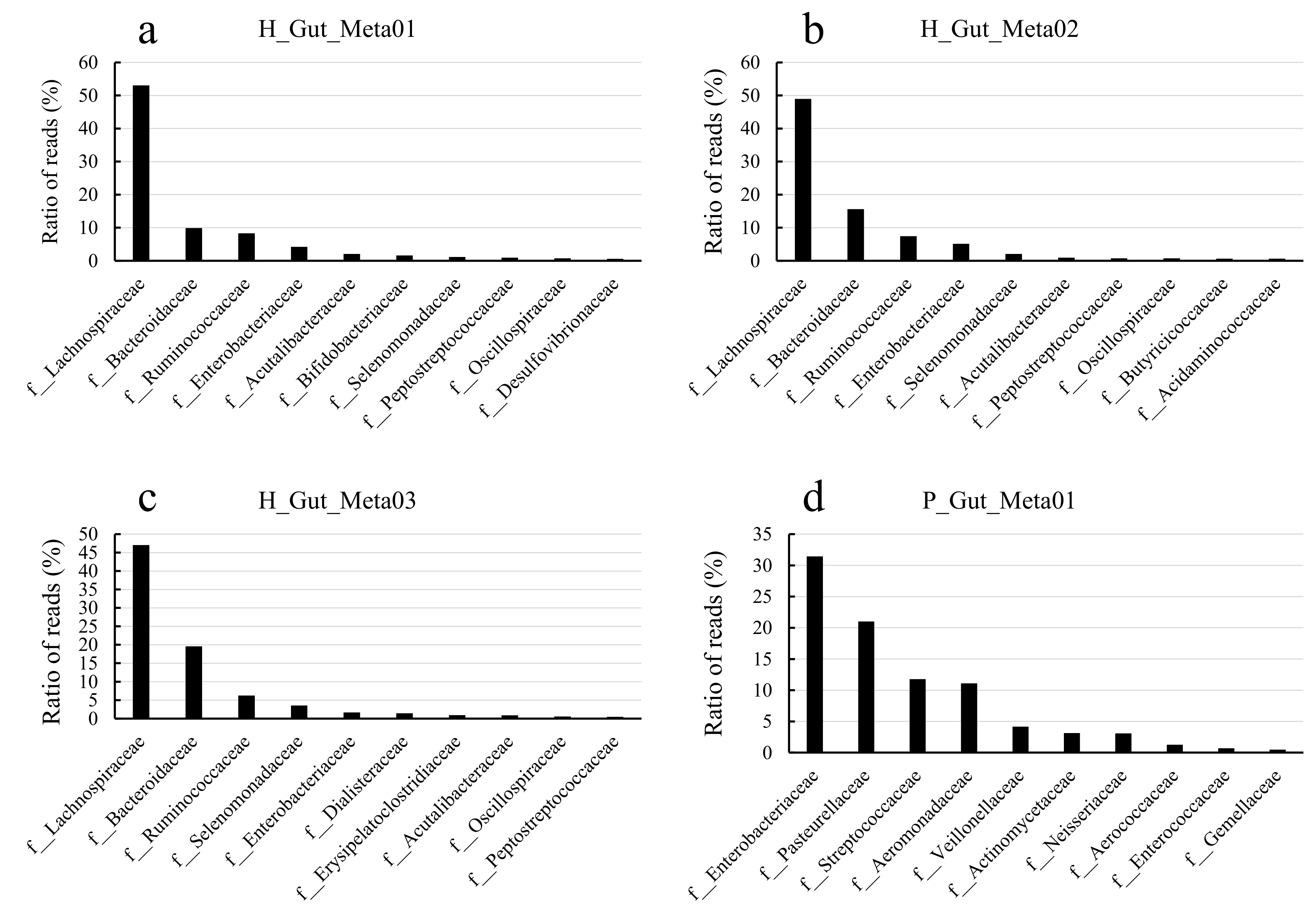

**Figure S6** Distributions of classified reads at different families for four gut samples.

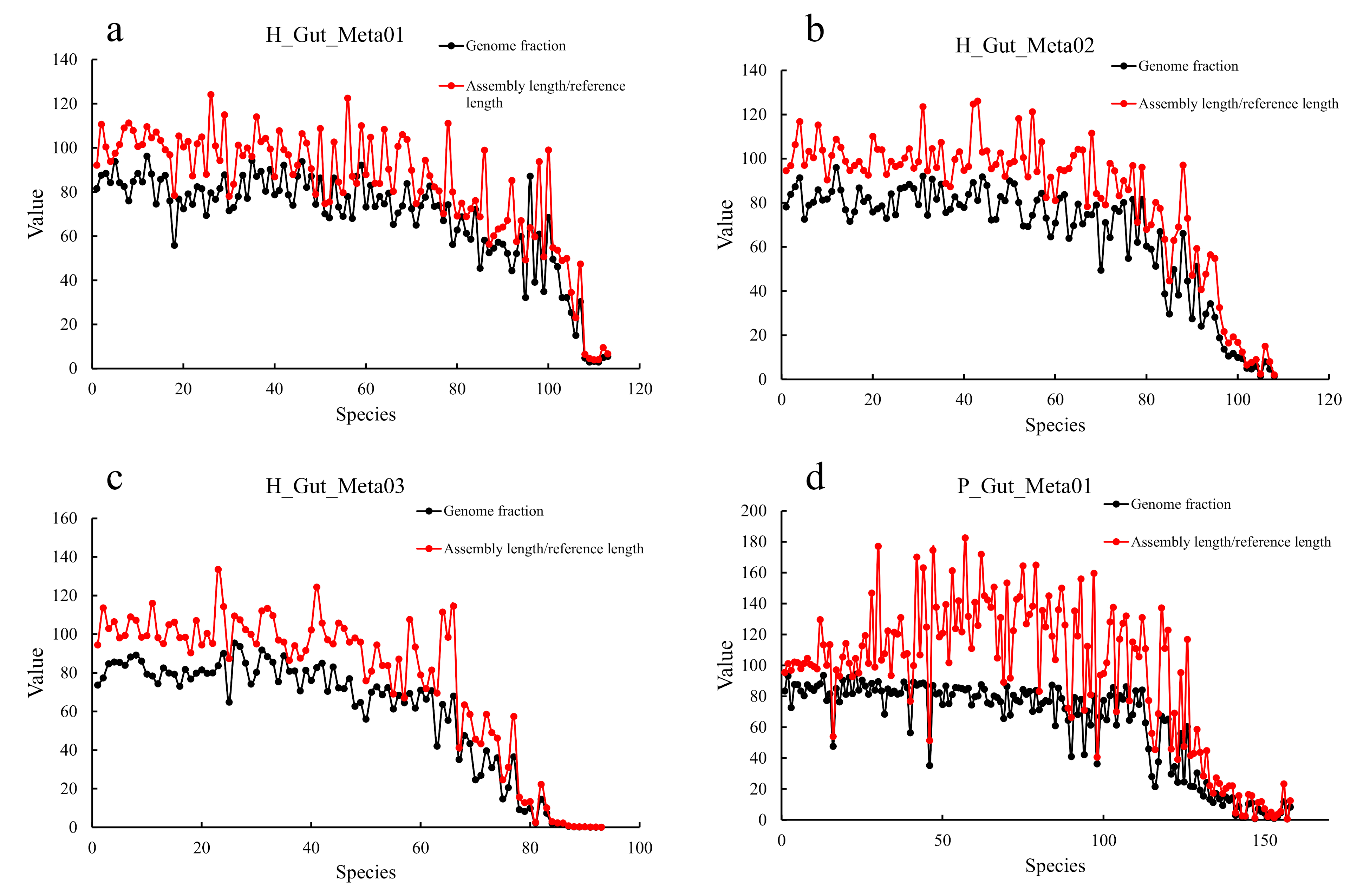

**Figure S7** Genome faction and ratio of assembly length to reference length of all species assembled in MetaTrass for four gut samples, and the species are ordered by the completeness.

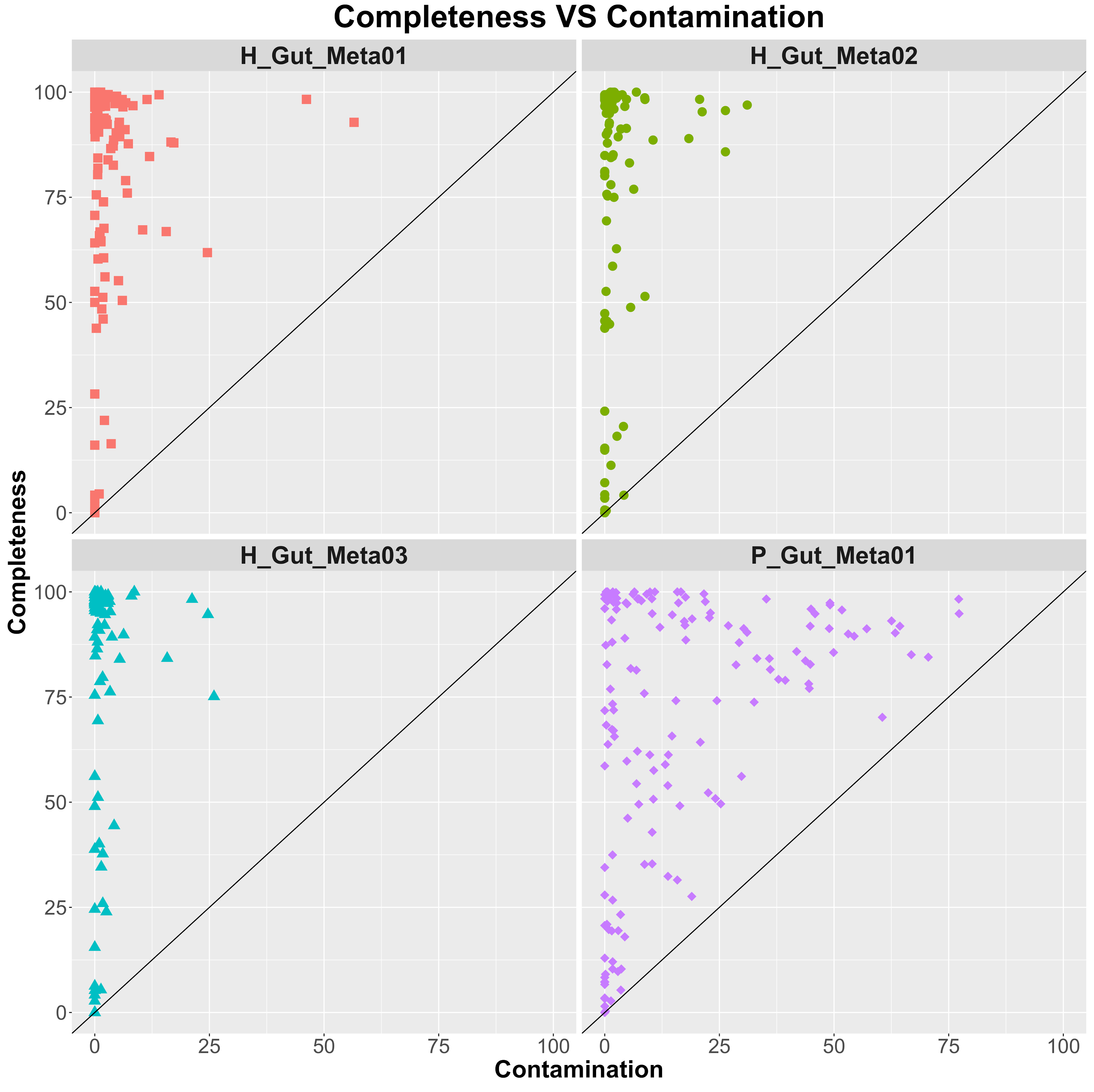

**Figure S8** Two-dimensional scatter plot of completeness and contamination evaluated by CheckM for four gut samples.

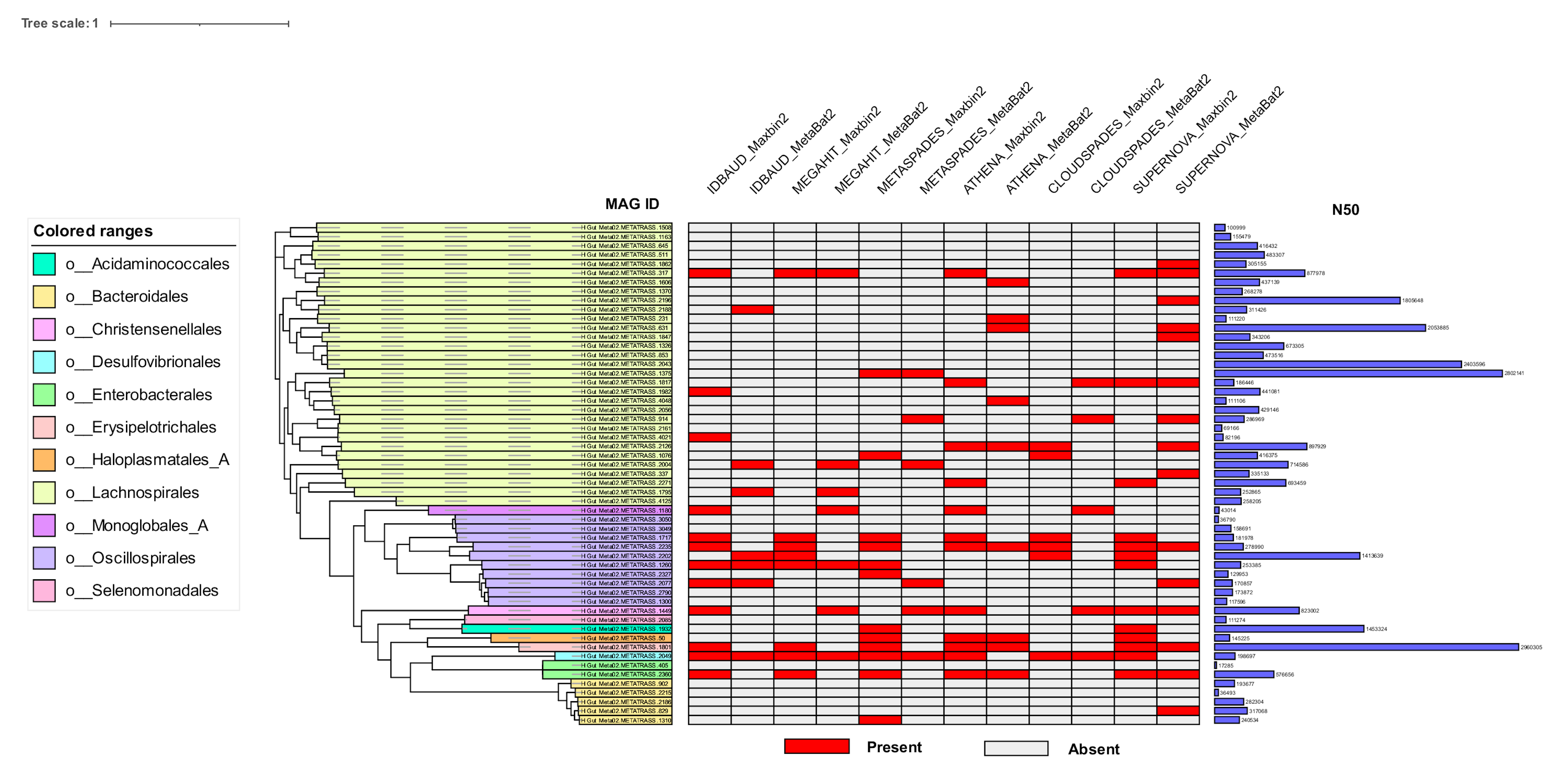

**Figure S9** Phylogenetic tree of the high-quality genomes assembled by MetaTrass for H_Gut_Meta02. The phylogenetic tree is on the left. Distribution of the high-quality genomes assembled by other methods are colored as red in the middle heat map. N50 of each high-quality genome is shown in the right histogram.

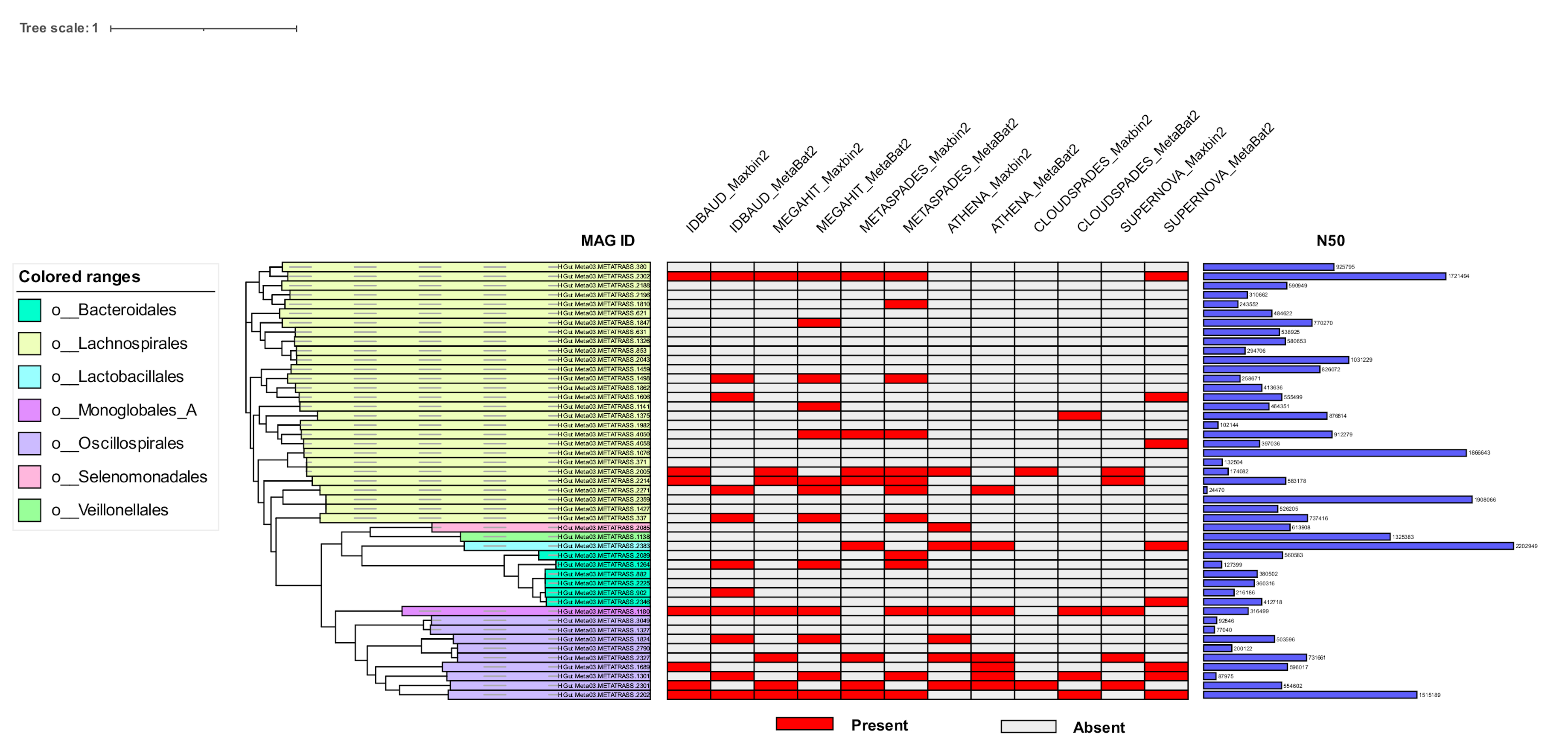

**Figure S10** Phylogenetic tree of the high-quality genomes assembled by MetaTrass for H_Gut_Meta03. The phylogenetic tree is on the left. Distribution of the high-quality genomes assembled by other methods are colored as red in the middle heat map. N50 of each high-quality genome is shown in the right histogram.

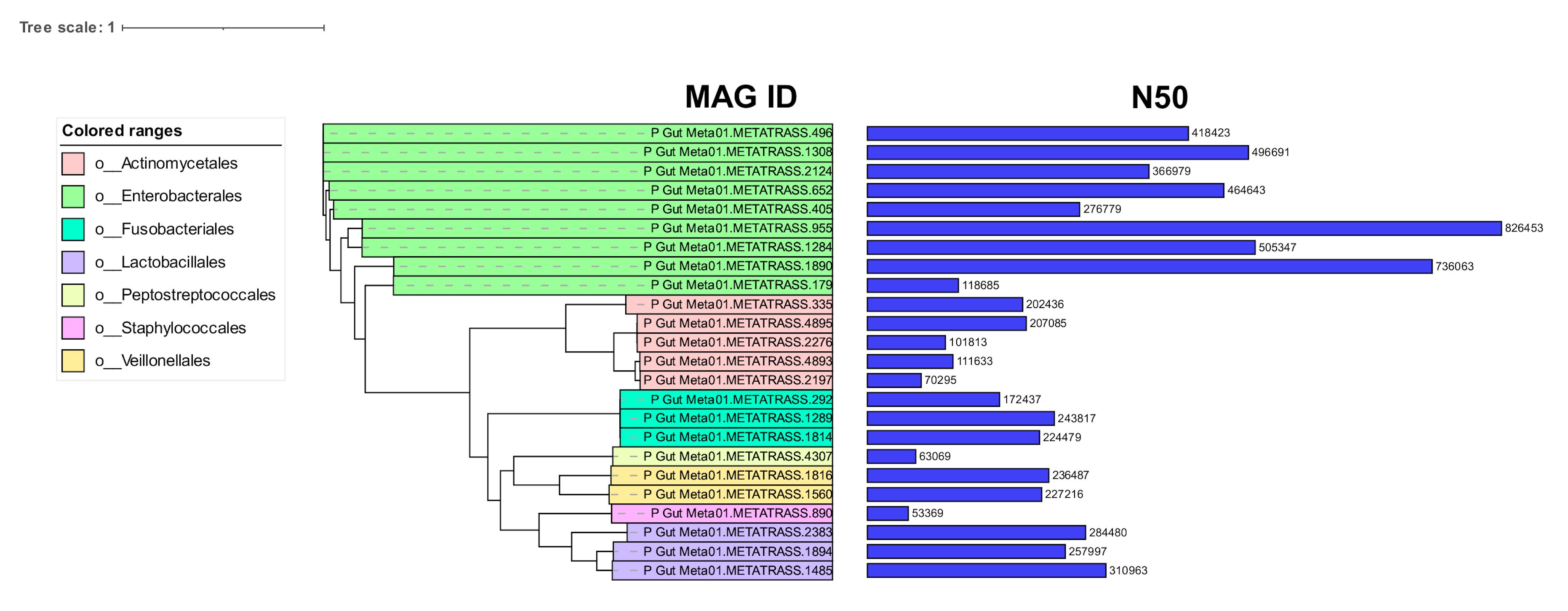

**Figure S11** Phylogenetic tree of the high-quality genomes assembled by MetaTrass for P_Gut_Meta01**.**  The phylogenetic tree is on the left. N50 of each high-quality genome is shown in the right histogram. Because the genome bins obtained by the combination strategies cannot be classified into species by GTDB-tk, the heat map is not showed for this sample.

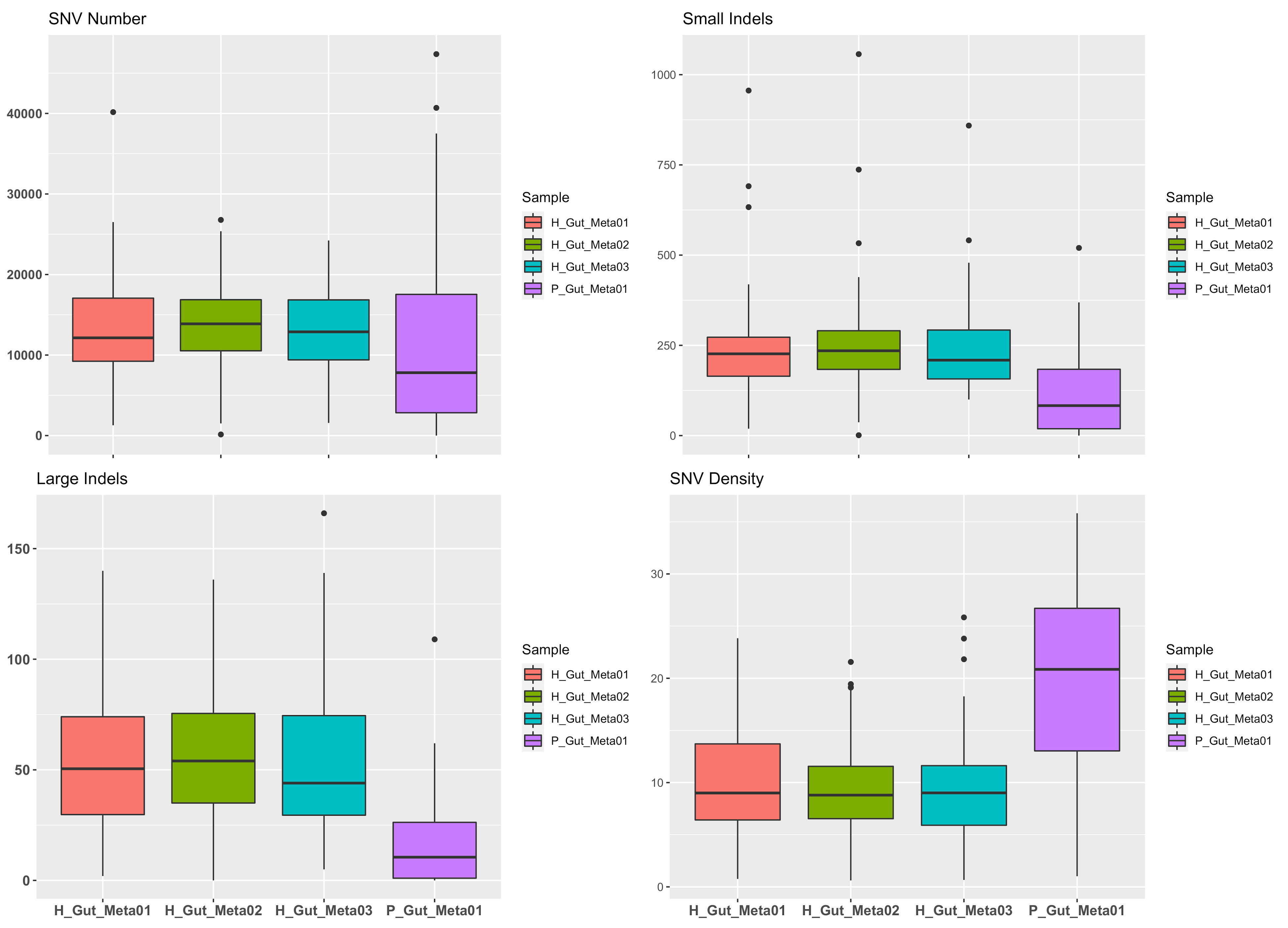

**Figure S12**  Box plot of variations. Box plots of SNVs (a), small indels (b), large indels (c) and SNV density (d) called from the high-quality genomes for four gut samples.

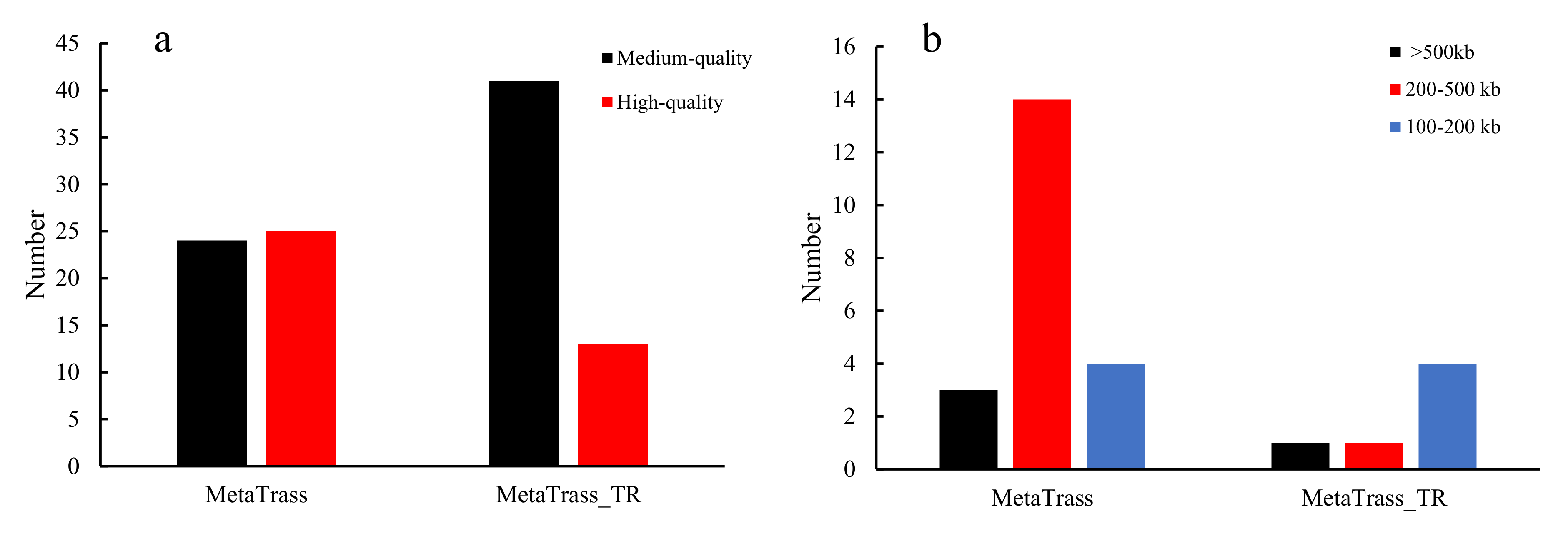

**Figure S13** Number of genomes with different quality (a) and contiguity (b) assembled by MetaTrass and MetaTrass_TR for the patient gut sample. Since MetaTrass_TR excluded the co-barcoding refining process compared to MetaTrass, the input dataset of co-barcoding assembling in MetaTrass_TR is the taxonomic reads set. Only the high-quality genomes are considered to count the number of genomes with different contiguity.
